# Structural flexibility dominates over binding strength for supramolecular crystallinity

**DOI:** 10.1101/2023.09.04.556250

**Authors:** Vincenzo Caroprese, Cem Tekin, Veronika Cencen, Majid Mosayebi, Tanniemola B. Liverpool, Derek N. Woolfson, Georg Fantner, Maartje M.C. Bastings

## Abstract

Supramolecular crystallinity is abundantly present in nature and results from directional, weak non-covalent interactions between components. Bottom-up nanotechnology aims to exploit such phenomena to control the self-assembly of ordered networks and complex objects from rationally designed monomers. Like all crystalline materials, 2D supramolecular crystals develop from an initial nucleation site, followed by growth, based on directional interactions. Traditionally, the binding strength and directionality of interactions is thought to dictate the nucleation and crystal growth, whereas structural flexibility favours defects. Usually, macromonomers present multiple binding units with relative intramolecular flexibility that affects their intermolecular interactions. Thus far, the effects of such flexibility on supramolecular assembly have not been explored. Here we introduce the concept of “interface flexibility” and demonstrate its critical importance in the nucleation and growth of supramolecular crystalline networks. We show that tuning the interface flexibility greatly expands the available design space for synthetic supramolecular crystalline materials.

## INTRODUCTION

Dynamic, yet controlled self-organisation of macromolecules into temporal geometric domains allows for the engineering of functional interfaces for catalysis, materials and nanotherapeutics.^1,2^ While challenging for engineers, these supramolecular crystalline networks are fundamental for many biological processes. For instance, the assembly of clathrin triskelions into polygonal patches enables cellular uptake^3–5^, and the organisation of TRIM5a in hexagonal patterns on the HIV-1 capsid combats viral infection.^6^ Traditionally, the main parameters used to design directional multivalent interactions are binding strength (affinity) and number of binding events (valency).^7^ Interestingly, both clathrin and TRIM5a use additional structural changes to transition from monomers into their characteristic geometric patterns.^3,6^ Consequently, the nucleation of these crystalline networks is influenced by local changes in mechanical properties of the monomers. These mechanical properties influence the spatial tolerance of the intermolecular binding interface and thereby impact the systems’ capacity to acquire long-range crystalline order. While the structure-function connection of these biological phenomena is evident, their complex multicomponent environment limits a systematic manipulation of molecular parameters to explore the role of interface mechanics in nucleation and growth of supramolecular crystalline networks.

Crystal formation is a complex process that involves both physical and chemical mechanisms, yet always starts with several molecules coming together to create a stable nucleation event.^8^ This process is driven by many properties of the system, balancing concentration, diffusion and surface energies.^9,10^ In traditional covalent crystals, the bonding between atoms is typically strong and directional, yielding a highly ordered and stable crystal structure.^11^ Supramolecular crystals often show a more flexible and dynamic structure, attributed to the weaker non-covalent interactions and larger macromolecular monomer unit-cell out of which they are built.^12,13^ This implies that structural mechanics of this macromolecular unit cell plays a critical role in the formation of supramolecular crystals.

Similar to nucleation in covalent crystals, non-covalent interactions in supramolecular crystalline materials have a directional component at the intermolecular interface.^14^ A deepened understanding of the flexibility limits of bond directionality at the supramolecular interface could translate into novel engineering strategies to tailor the nucleation of such materials. To enable a detailed molecular analysis of the structural design parameters that influence the nucleation and growth mechanisms of supramolecular networks, modular monomers without complexity of biology are essential. While protein design and engineering approaches toward geometric networks have shown beautiful lab-made examples^15–18^, modulation of flexibility and affinity is not straightforward without altering the monomer building block. At micron length scales, colloidal supramolecular crystals guided by molecular recognition at the interface present impressive degrees of programmable design,^19–21^ yet the effect of nuanced changes in structural mechanics get lost in the overall system dimensions.

Compared to protein-based monomers and colloids, DNA-nanotechnology^22^ offers a predictable and modular platform to decouple the interplay between macromolecular flexibility and monomer-monomer affinity. When using synthetic DNA as material tool, structural flexibility can be controlled by the balance of single-strand (ss) versus double-strand (ds) sections, as well as the overall length compared to the persistence length of the DNA double helix.^23^ Affinity can be manipulated through the length of single strand complementary ends (so called “sticky ends”) or the nucleotide sequence at the end of a “blunt end” double helix, as GC presents a stronger *π*-*π* stacking interaction than TA.^24^ Utilizing the multivalency concept, a higher affinity interaction can be designed by systematic multimerization of adjacent DNA blunt-ends. Combined, these characteristics make DNA an ideal model material to explore towards what extend rigidity at the intermolecular interface guides the nucleation-growth mechanisms in supramolecular 2D crystalline materials.

## RESULTS AND DISCUSSION

### Minimal structural changes lead to different network architectures

DNA-based macromonomers assembled from a limited number of oligonucleotides (“tiles”) have been designed and used to engineer a range of simple to highly complex supramolecular systems.^25^ We used the established DNA 3-point-star (3PS) motif as starting point, which allowed us to build upon existing work on the supramolecular crystalline self-organisation of DNA-tiles^26^ (Supplementary Fig. 1). This macromolecule presents a 3-fold rotational symmetry and consists of seven ssDNA strands - three unique sequences - organised in three four-arm junctions (Fig 1a). The conventional approach to obtain supramolecular crystals from DNA tiles is via the display of short ssDNA terminal ends that enable so-called “sticky-end” hybridization with their sequence-complement. These networks assemble in solution and show long-range order with very few defects.^27^ However, the use of sticky-ends comes at the cost of kinetic control and renders the analysis of self-assembly mechanisms related to monomer design obsolete. In a blunt-ended variant, the interaction-interface consists of a multivalent array of *π*-*π* stacking units, presented by the nucleobases. When multiple DNA helices are present in parallel, the distance from a DNA cross-over junction affects the spatial freedom at the end of a DNA helix extension.^28^ The geometric alignment of multiple of these weak non-covalent interactions allows for a sufficiently strong directional force to self-assemble into 2D crystalline networks. However, they are also reversible, and their inherent dynamics permit targeting the nucleation-growth mechanisms of network formation.^29^

**Fig. 1.**
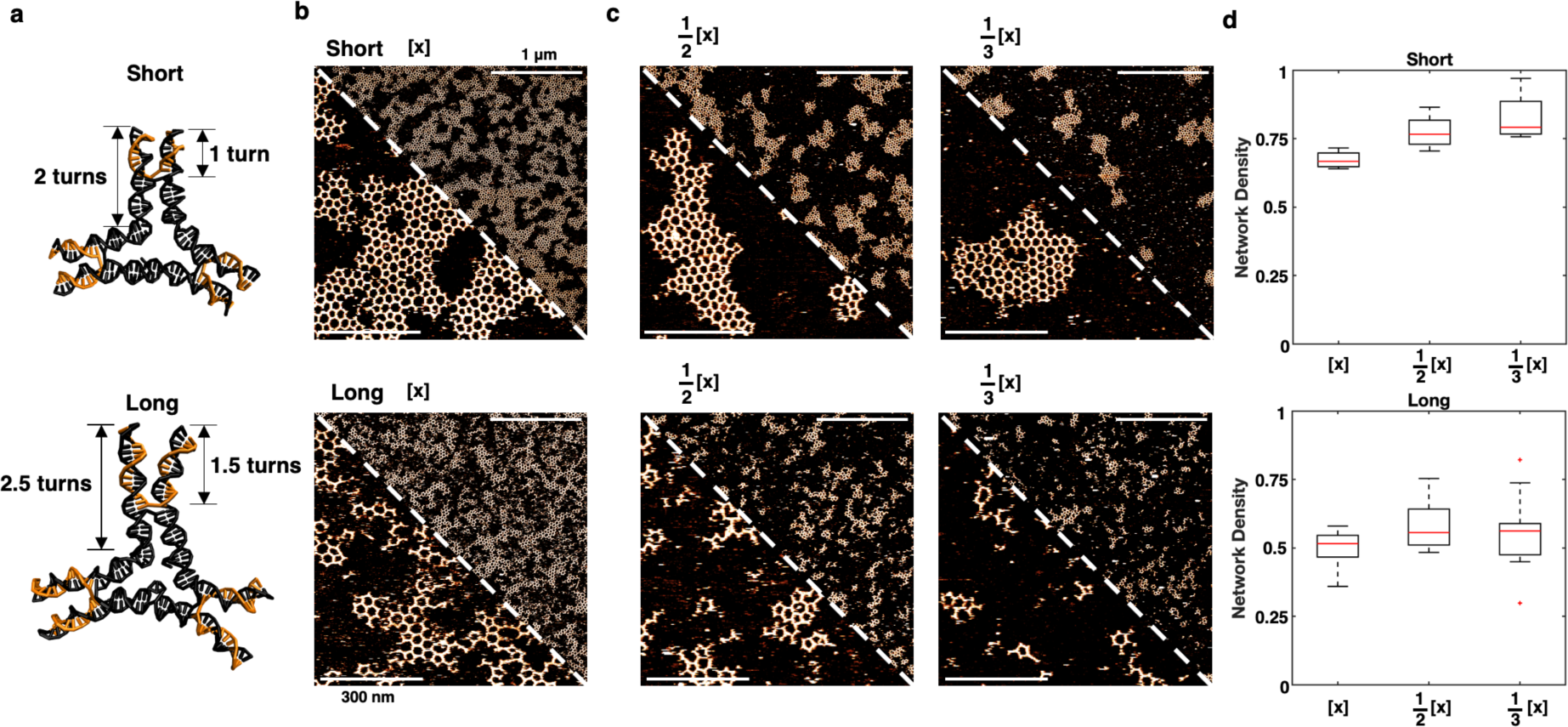
Steady-state assembly of hexagonal lattices formed by *short* and *long* 3PS. **a,** DNA blueprint of the *short* and *long* monomers obtained from Molecular Dynamics simulations, indicating the overall and terminal arm-lengths in double-helical turns. **b,** AFM images of the steady-state networks formed by the *short* (top row) or *long* (bottom) monomers. **c,** AFM images of the steady-state networks formed by the *short* (top row) or *long* (bottom) monomers at decreasing concentration. **d,** Weighted mean of the Network Density as a function of monomer concentration. The red lines indicate the median. Outliers, defined as data points beyond 1.5 times the interquartile range, are represented as individual points. Upper scale bars 1 µm, zoomed-in lower scale bars 300 nm. Conditions: Overnight incubation on mica at room temperature in the presence of 10mM MgAc_2_. [x] is 6nM for *short* and 3.8nM for *long* 3PS.

To explore the impact of monomer mechanics on the growth and lattice structure, we made two 3PS variants: one with arms of 2 DNA turns (Fig. 1a, “*short*”); and a second 3PS with 0.5-turn extensions per arm (Fig. 1a, “*long*”). The modification generates a larger star motif in terms of absolute size and corresponding global flexibility, but also the spatial tolerance of the arm termini (terminal flexibility) is increased. This terminal flexibility results from the distance after the Holliday junction in the middle of the 3PS arm (Fig. 1a), allowing the two parallel double-helices to have more freedom and move independently. We let both systems assemble overnight in aqueous buffer solution on mica (Supplementary Fig. 2) and imaged the steady-state networks by atomic force microscopy (AFM). Clear differences between the steady-state assemblies of the *short* and *long* 3PS were observed (Fig. 1b). The *short* monomers predominantly formed large hexagonal networks, whereas the *long* monomers assembled into smaller, elongated islands. To explore if the difference between the designs was related to the monomer concentration, we imaged both systems over a range of concentrations (Fig. 1c). Measurements at higher concentrations led to surface crowding and multilayer assembly (Supplementary Fig. 3). For all dilutions, we consistently observed large islands for *short* 3PS and many small structures for *long* 3PS.

We developed a tailored detection algorithm in MATLAB^30^ to identify each island, 3PS monomer centres, and polygons present in a frame reliably (Methods, Supplementary Fig. 4). By considering each island as a network of polygons, we applied the concept of network density (*ND*)^31^ to quantify the degree of ideal radial growth within the assembled islands. For our case, *ND* was expressed as the ratio between the number of observed polygons in an island (*R_o_*) and the number of polygons in an ideal island formed by the same number of monomers arranged in a hexagonal symmetry and assembled by radial growth (*R_i_*) (Supplementary Fig. 5). Thus, *ND* values close to 1 indicate observed islands that are close to an ideal crystalline network. As *ND* drops, the monomers are increasingly less organized in this way, but for instance, as more elongated assemblies. Application of the *ND* to all steady-state results confirmed the presence of 2 distinct supramolecular network architectures (Fig 1d.). The *short* 3PS follow the assembly of a radial crystalline network much closer than the *long* variant. *ND* approaches 1 for the *short* 3PS measured at the lowest concentration. We deduce from this that radial growth is hindered by coalescence of multiple islands at the higher concentrations. By contrast, the low *ND* value characteristic of the elongated assemblies seen for the *long* 3PS is concentration independent. Thus, the small change in arm-geometry and associated terminal flexibility has a significant impact on the global network self-assembly.

### Rigidity and affinity cooperate for a crystalline organization

Next, we considered if changes in intermolecular binding affinities could overcome the negative effects on lattice propagation caused by structural flexibility. For DNA-based macromonomers, these affinities can be tuned by the number of parallel double-helices (two in each 3PS arm) or the sequence of terminal nucleotides participating in the blunt-end *π*-*π* stacking interactions.^24^ When fully surrounded by neighbours in a hexagonal arrangement, 3×2 directional *π*-*π* interactions are formed, stabilizing otherwise weak binding through the strength-in-numbers principle of multivalency. Since the *π*-*π* stacking energy is defined by the chemical structure of the DNA nucleobase, the overall binding affinity between monomers can be controlled through the sequence of the terminal nucleotides. Therefore, we modified the original *long* and *short* 3PS motifs, which had GC-TA as terminal nucleotide pairs, to present either TA-TA or GC-GC (Fig. 2a, Supplementary Fig. 6-7). These modifications directly affect the *π*-*π* stacking energy, expected to range from ΔG ∼2 kcal mol^−1^ for TA-TA, to ΔG ∼6.8 kcal mol^−1^ for GC-GC.^24^ The strain within our supramolecular network is expected to be minimal when all monomers arrange into a hexagonal array, as all arms of the 3PSs are in a relaxed orientation, maximizing the number of *π*-*π* binding events. However, due to the global flexibility present in the monomer structure, different polygons can be formed (so-called “defects”), which can be corrected by breakage of the *π*-*π* interactions and reorganization of the macromolecules into the ideal hexagonal lattice structure; *i.e.,* the use of weak non-covalent interactions allows for reversibility and subsequent optimization (annealing) of the system. We hypothesized that besides affecting global network formation, the choice of terminal nucleotide identity might also impact the local polygonal composition.

**Fig. 2.**
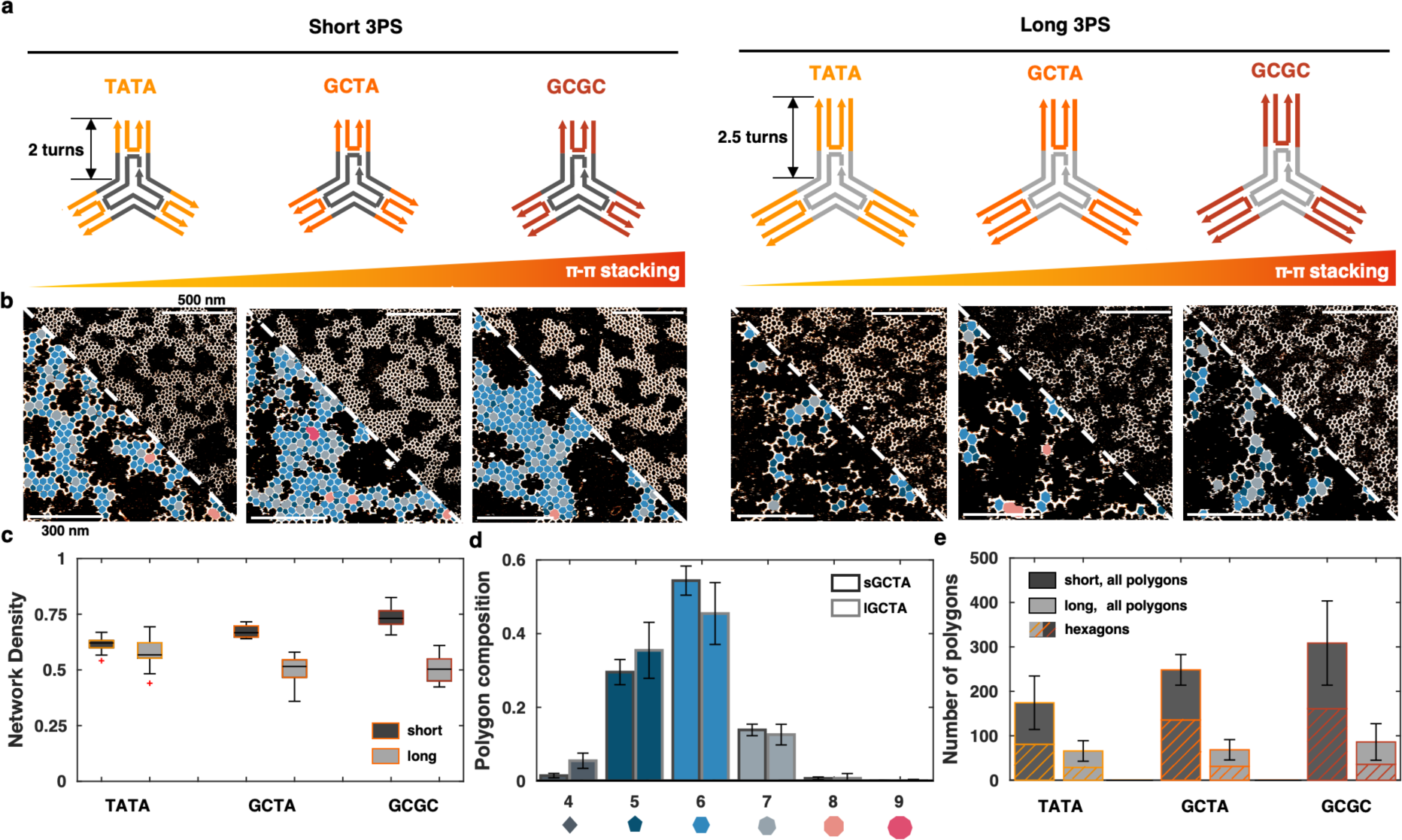
Quantifying the effect of intermolecular affinity on network crystallinity. **a,** 3PS library, where each monomer has a unique combination of an arm length (2 or 2.5 turns) and a terminal base-pair (TA-TA, GC-TA or GC-GC). The blunt-end *π*-*π* interaction strength increases from left to right (from TA-TA to GC-GC). **b**, (top right triangle) AFM images of the assembly in steady-state (after 24 h) for each 3PS design. (bottom left triangle) Higher magnification and overlay of the polygon identification algorithm (pentagons, dark blue; hexagons, blue; heptagons, light gray). **c,** ND of all 3PS assemblies showcasing a moderate effect of affinity in short 3PS systems yet marginal impact in long 3PS systems. The boxes represent the middle 50% of the data, with the bottom and top of the box representing the 25th and 75th percentiles, respectively. The red lines indicate the median. Outliers, defined as data points beyond 1.5 times the interquartile range, are represented as individual points **d,** Polygon distributions for short GCTA versus long GCTA assemblies, using at least 12 regions with an area of 750 x 750 nm in at least 2 independent experiments. **e,** Average count of all polygons, with the hexagon contribution highlighted by the striped surface. Error bars represent the standard deviation.

The six 3PSs variants were assembled separately overnight and subsequent steady-state AFM showed consistent patterns within the groups of *short* and *long* monomers, (Fig. 2b), suggesting that changes in intermonomer affinity had little or no impact on the overall network structure. From the *ND* values, increasing the inter-monomer affinity for *short* 3PSs slightly enhanced the radial crystal organization, whereas no clear effect was apparent for the *long* 3PSs (Fig. 2c). Interestingly, the weakest *short* 3PS (TATA) approaches the organization of the *long* 3PSs, yet the strongest *long* 3PS (GCGC) does not approach that of the *short* 3PS system. Thus, within the range of interactions present in these 6 variants, mechanical properties (affecting the entropy of the system) apparently dominate the interaction affinity (affecting the enthalpy balance) in terms of network formation.

For a quantitative analysis of the internal polygon composition, we extended our image-processing algorithm to automate the detection and labelling of polygonal geometries from 3 to 9 monomers in *short* GCTA versus *long* GCTA assemblies (Fig. 2d; same analyses of TATA and GCGC are in Supplementary Fig. 8). This revealed that hexagonal organization is prevalent, with pentagonal and heptagonal assemblies responsible for the majority of “defects”. Similar heterogeneity in polygon distribution has been observed in clathrin networks and synthetic peptide architectures, and can be explained by a moderate structural flexibility within the monomers.^3,4,32^ Plotting the hexagonal content within all polygons leads to two conclusions (Fig. 2e): First, the total amount of assembled polygons is significantly higher for the short compared to the *long* 3PS molecules, indicating a favoured network growth mechanism. Second, the hexagonal content within the *short* 3PS is moderately dependent on the terminal affinity, with short GCGC and short GCTA forming the most crystalline network, as judged by the fewest polygonal defects and highest ND (Fig. 2e). This suggests that rigidity and affinity cooperate to create a more-crystalline organization and that there is an affinity threshold after which hexagonal order is favoured.

### Real-time imaging of network formation reveals differences in self-assembly mechanisms

The steady-state data for the 3PS variants suggest fundamental differences in the assembly mechanism between the short and long monomers. Unfortunately, static AFM cannot provide details on the real-time dynamics of network formation. Therefore, we turned to high-speed AFM (HS-AFM)^33–35^ in photothermal off-resonance tapping (PORT) mode^35,36^ in an attempt to visualize and analyse the mechanisms of assembly from the earliest interactions. With a frame rate of 0.4 fps, the organization of 3PS macromolecules into the typical hexagonal patterns was observed from the moment of monomer injection (Fig. 3a, Supplementary Videos 1-6). This enabled us to focus on the initial phases of supramolecular crystal nucleation.

**Fig. 3.**
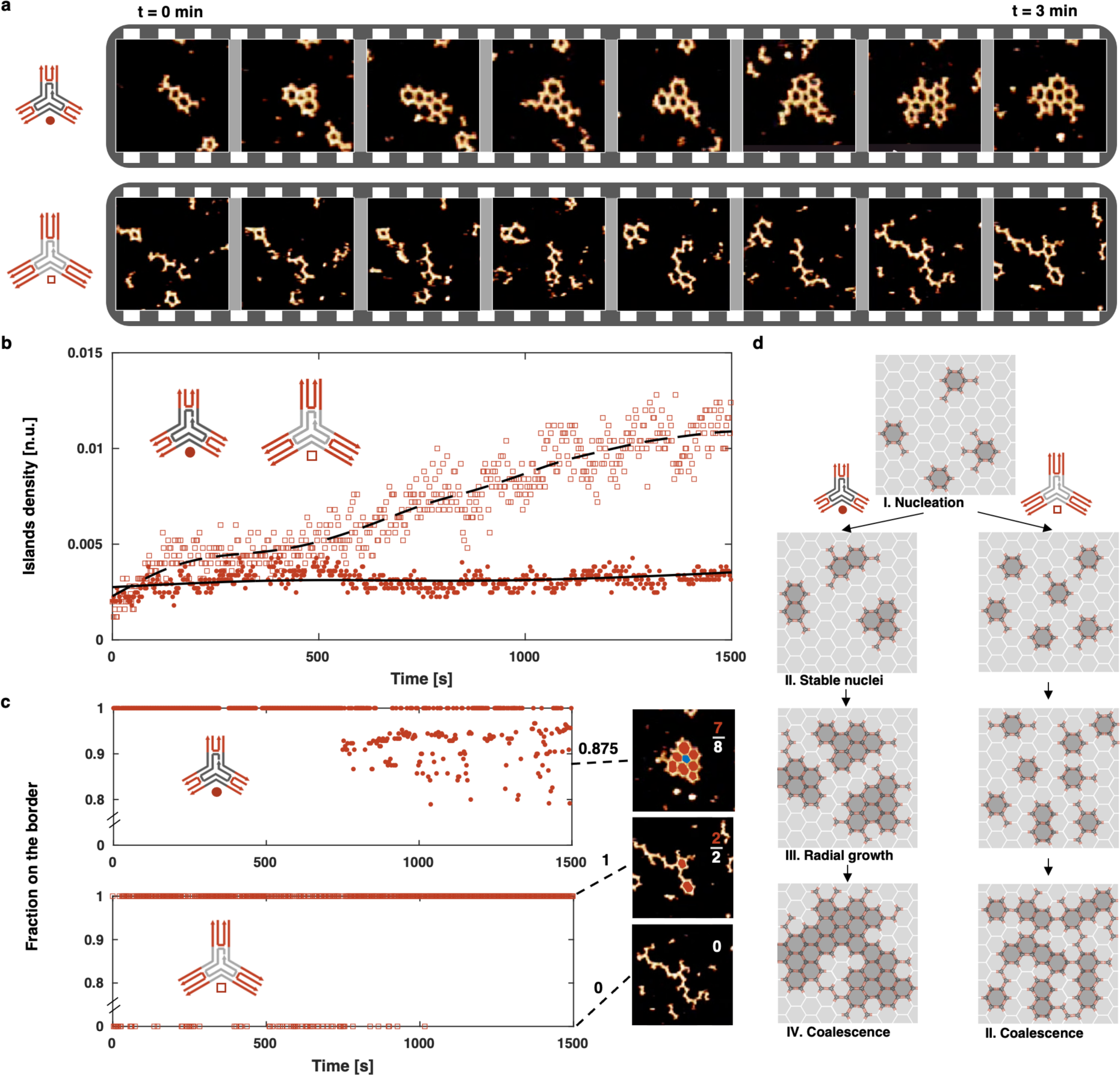
HS-AFM detection of 2 distinct growth mechanisms. **a,** Time-lapse of *short* (top) and *long* (bottom) 3PS self-assembly over 3 minutes highlighting the difference in nucleation. Snapshots taken from full length timeseries. **b,** Detected *island density* over time (25 minutes imaging) using tailored particle tracking algorithms for *long* and *short* 3PS. **c,** Quantification of radial crystal island growth through detection of fraction of polygons on a border (coloured red) over the total number of polygons. When no polygons are present, the ratio is taken to be 0. **d**, Schematic representation of the nucleation-growth self-assembly mechanism of *short* 3PS and coalescence-based elongated island formation of *long* 3PS.

Even on qualitative inspection, it was immediately apparent that the *short* and *long* 3PS monomers had different self-assembly pathways (Fig. 3a): whereas the short monomers quickly formed stable closed networks, the *long* 3PS were highly dynamic and seemed not to enter a stable growth regime. The *short* 3PSs formed a small number of large islands indicative of stable initial nucleation events followed by radial growth. By contrast, multiple, small, and elongated islands observed in the *long* 3PS samples suggest a lack of stable nucleation. To quantify these differences, we identified all islands and particles present over time using the aforementioned image analysis (Fig. 3b, Supplementary Fig. 4). The density of islands remained constant for the *short* 3PS, whereas for *long* 3PS the number of islands grew, all while the density of monomers on the surface steadily increases due to the continuous surface adsorption of monomers from the bulk (Supplementary Fig. 9). This confirms that incoming monomers attach to a stable nucleus and contribute to the growth of the supramolecular crystal for the *short* 3PS; whereas incoming *long* 3PS monomers do not nucleate and eventually crowd the surface and coalesce into larger elongated structures.

We next counted the number of polygons present on the border of the biggest island and expressed it as a ratio in respect of the total number of polygons in that island (Fig. 3c). When the network grows, an increasing number of polygons will be in the centre of an island (Fig. 3c, blue), causing a decrease of this *border ratio*. Following an initial nucleation time, which for the *short* 3PS systems was ≈10 minutes, the first centre-monomers (e.g. *border-ratio* <1) started to appear. Subsequently, the border fraction further decreased, which is a strong confirmation of radial island growth from an initial stable nucleus. The *long* 3PS showed a very different profile as during the time of the experiment, all monomers remained present at a border, either as polygon (“1”) or in an elongated polymeric conformation (“0”).

Combining HS-AFM and the tailored particle-detection analyses, we propose the following kinetic pathways for the assembly of the two 3PS systems, Fig. 3d. Considering the *short* 3PS system first, as *π*-*π* stacking is a weak interaction, the blunt-ended 3PS motifs only interact laterally when on a surface where they can diffuse freely as monomers. Due to the parallel alignment of DNA double-helices in the arms of the 3PS motifs, a multivalent array of *π*-*π* interactions is presented. Together with the structural tri-symmetry, stability of the assembly increases by multimerization of the *π*-*π* stacking interactions leading to the formation of polygonal nuclei. These nuclei are less diffusive than the 3PS monomers, because the nuclei have larger contact areas and more interactions with the mica surface.^37^ This reduced mobility allows the formation of stable nuclei and subsequent growth of supramolecular crystals. By contrast, for the *long* 3PS, visibly no stable nuclei are formed, which prevents the transition to a radial growth phase. Eventually, however, as the density of monomers at the surface increases, the many small nuclei coalesce to create elongated-island structures (Fig. 3d) as observed in steady-state analysis.

### Interface flexibility is the determinant for stable nucleus formation

With two distinct kinetic pathways evident, next, we explored the main molecular features that might cause this change in assembly mode. The observed difference in stable-nucleus formation suggested the interface between the two types of monomer to be significantly different. These interfaces are formed by a pair of double-stranded DNA helices that make up each arm of the monomers. Considering the strong directional nature of the π-π stacking, the strength of interaction between two different particles is directly affected by the possible orientations that these helices (and in particular their termini) can assume: it will be maximised when the two helices are precisely parallel and aligned, and fall off as the termini are oriented in different directions, to a breaking point where just one weak π-π bond can form. This lets us define two possible extreme states for an arm: “open”, if the double-helices are point in different directions; and “closed”, if sufficiently parallel to maximise π-π interactions (Fig 4a). To assess the possible distributions of *short* and *long* 3PS in these states, we used oxDNA^38,39^ (Supplementary Figure 10-11) and a series of Molecular Dynamics^40^ simulations (Supplementary Methods). The sampled probability distribution functions of the end-to-end distance between DNA helices in an arm (Fig. 4b) show that the *short* 3PS presents a narrower distribution (∼2 nm, centred around ∼2.7 nm) compared with that for the *long* 3PS (∼ 6 nm, centred at ∼3.2 nm). Comparing the distributions to the spatial tolerance to form two simultaneous *π*-*π* stacking interactions, expected to be 1.5 – 2.5 nm, the arms of *short* 3PS are almost four times more likely to be oriented in a bond-forming conformation than the arms of *long* 3PS. Therefore, the chance that *long* 3PS makes productive interactions leading to stable nuclei is significantly reduced compared to the *short* 3PS units. This change in interface flexibility provides a potential underlying explanation for the different mechanisms of supramolecular assembly.

**Fig. 4.**
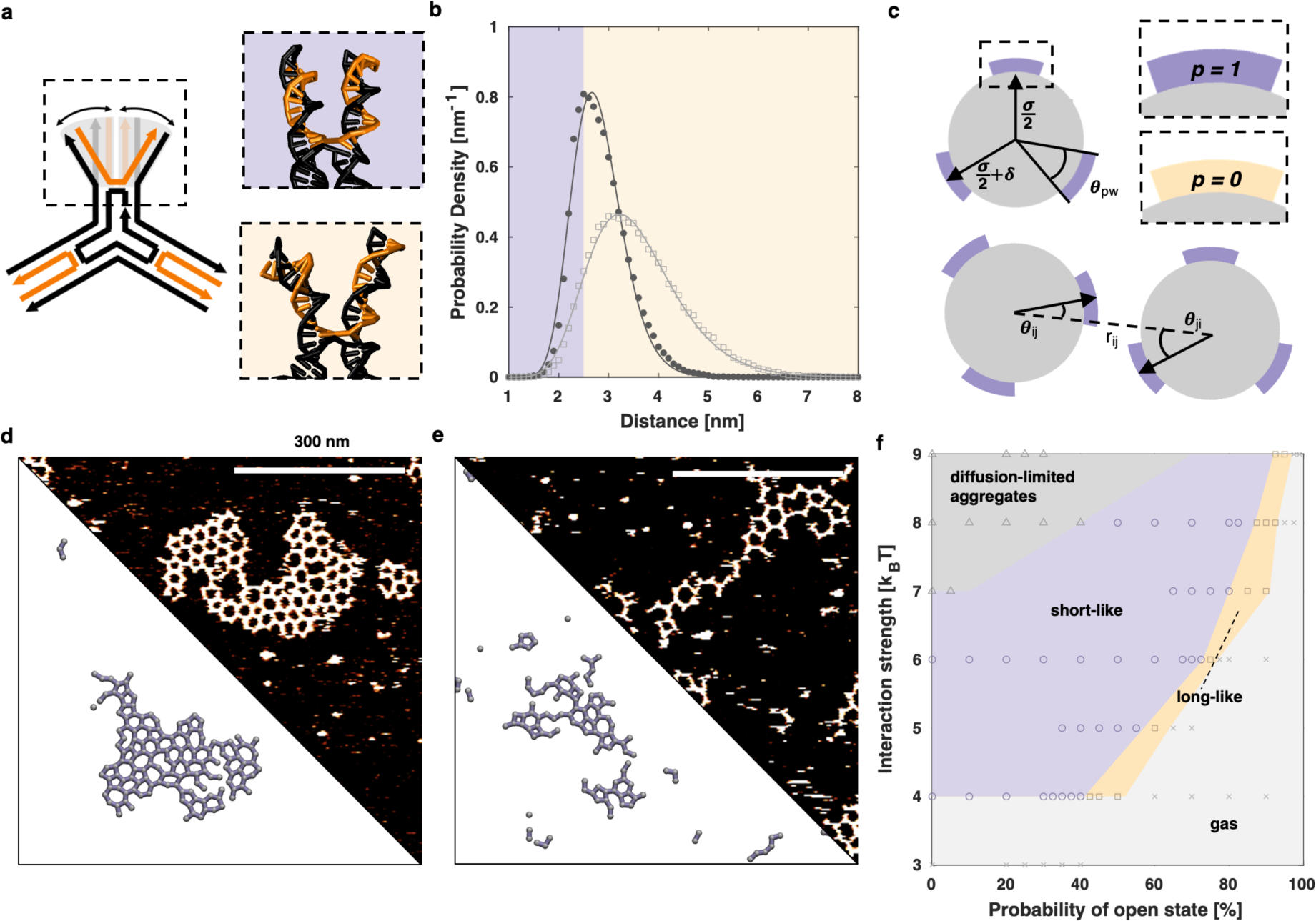
Importance of interface flexibility. **a,** Representation of interface flexibility related to the mobility of the peripheral segments of DNA following the Holiday junction. The snapshot in the upper box represents an aligned, “closed” configuration that allows directional binding; whereas the snapshot in the lower box represents a wider, “open” configuration that is unavailable for binding. **b,** Probability distribution of the end-to-end arm distances at the π-π interface obtained from the oxDNA MD simulations of the 3PS monomers. The purple area highlights the region where the arms are in a closed state, whereas yellow is used for the open state. **c,** Illustration of a patchy particle model used for Monte Carlo simulations, highlighting the patch state with variable *p* (*p*=1 for closed state and *p*=0 for open state), and the *P*_*open*_ the probability that a patch has a state *p*=0. Other model parameters defined as in the classical Kern-Frenkel model, where *σ* is the diameter of the hard-core repulsion, *δ* is the attraction range, *θ_pw_*is the patch width. For more details, see SI. **d, e,** Comparison of simulation results, bottom, and steady state measurements, top of *short* 3PS (**d**) and *long* 3PS (**e**). The selected frames are taken from simulation trajectories with similar average ND values as in the experiments. **f,** Approximate phase diagram as a function of the probability of open state (i.e. when *p*=0) and the interaction strength. Dark grey region shows the diffusion-limited aggregation regime, purple/orange shows the short-like/long-like regime, and the light gray shows the gas (mostly single particles) phase. Each datapoint represents a set of conditions simulated, with length dimensions normalized to our *short* design. The phase diagram was obtained for *σ=1*, *δ=0.038* and *θ_pw_*=0.40.

We turned to Monte Carlo simulations^40^ to explore the effect of interface flexibility and spatial tolerance of the *π*-*π* interface, and their impact on supramolecular crystal nucleation in greater detail. A patchy-particle model was used to describe the 3PS motif as a particle made up of a repulsive core and 3 attractive sites equally distributed on its perimeter (Fig. 4c). This is a common strategy used for the study of directional interactions in macromolecular systems.^41–44^ The aforementioned “closed” and “open” states were introduced into a classical Kern-Frenkel^45^ potential through the concept of the “state” of a patch, where the state variable assumes a value of 1 when representing closed arms (indicating that the interface is available for binding) or 0 when the interface is open (detailed information about the model in Supplementary Methods). Similar to an Ising model^46,47^, each interface can switch states through a dedicated move following preassigned probabilities. We explored the parameter space representing the ensemble of interface flexibility by varying the state-switching probabilities and the strength of the interaction. This allowed us to recover states that are qualitatively and quantitatively similar to those obtained in our steady-state AFM measurements (Fig. 4d&e), confirming that the interface flexibility alone can define the nucleation-growth mechanism. Finally, using the *ND* and island size as observables to distinguish between different assembly mechanisms, we constructed a complete phase diagram (Fig 4f). We were able to observe states that match our *long* 3PS (anisotropic elongated clusters) and *short* 3PS, (compact isotropic clusters) labelled *long*-like and *short*-like, respectively. Additionally, we found conditions where only single molecules or small aggregates are present (similar to a two dimensional “gas”), which conform to literature reports.^48^ We also found that above a minimum interaction strength, there is always a range of opening probabilities for which anisotropic aggregates (the long-like state) were observed, even if they form reversible bonds. To the best of our knowledge, this is a unique case, since elongated structures are typically observed only when approaching the Diffusion Limited Aggregation conditions (i.e., in the limit of infinite interaction strength).^49–51^

### Restoring the interface rigidity returns stable nucleation

The clear change in self-assembly mechanism caused by differences in interface flexibility prompted us to redesign the *long* 3PS with the aim of installing a more-rigid interface like that of the *short* 3PS. For the initial *long* 3PS design, the arms were terminally extended by 5 nucleotides while the central DNA strand, and with it the cross-over point in the arms, were the same as for the short design (see Figs. 1a and 2a). As such, 2 additional sites of flexibility are present in the *long* 3PS design: a global arm flexibility due to the extended overall length; and the prolonged distance from the cross-over point results in a considerable local flexibility^28^ at the 3PS periphery (Fig. 5a,b). To reduce the interface flexibility and attempt to introduce a nucleation-growth profile for a *long* 3PS design, we moved the position of the cross-over point to make it similar to that the *short* 3PS design. This gave a new “*long-rigid*” (LR) 3PS design (Fig. 5a,b, Supplementary Fig. 12). Moving the DNA cross-over position in this way does not alter the chemical composition or overall monomer design, but it should change structural rigidity, exemplary of the strategic modularity present within DNA nanotechnology.^28^ Indeed, OxDNA simulations of the three 3PS designs (*short*, *long*, and *long-rigid*) showed changes in global and peripheral flexibility, and confirmed that by moving the cross-over position, the long-rigid 3PS monomer had a local interface rigidity similar to the *short* 3PS (Fig. 5b, Supplementary Fig. 13). Since the overall arm length is the same as the original *long* 3PS, the global flexibility resulting from the overall arm length corresponds to that found in the original *long* 3PS.

**Fig. 5.**
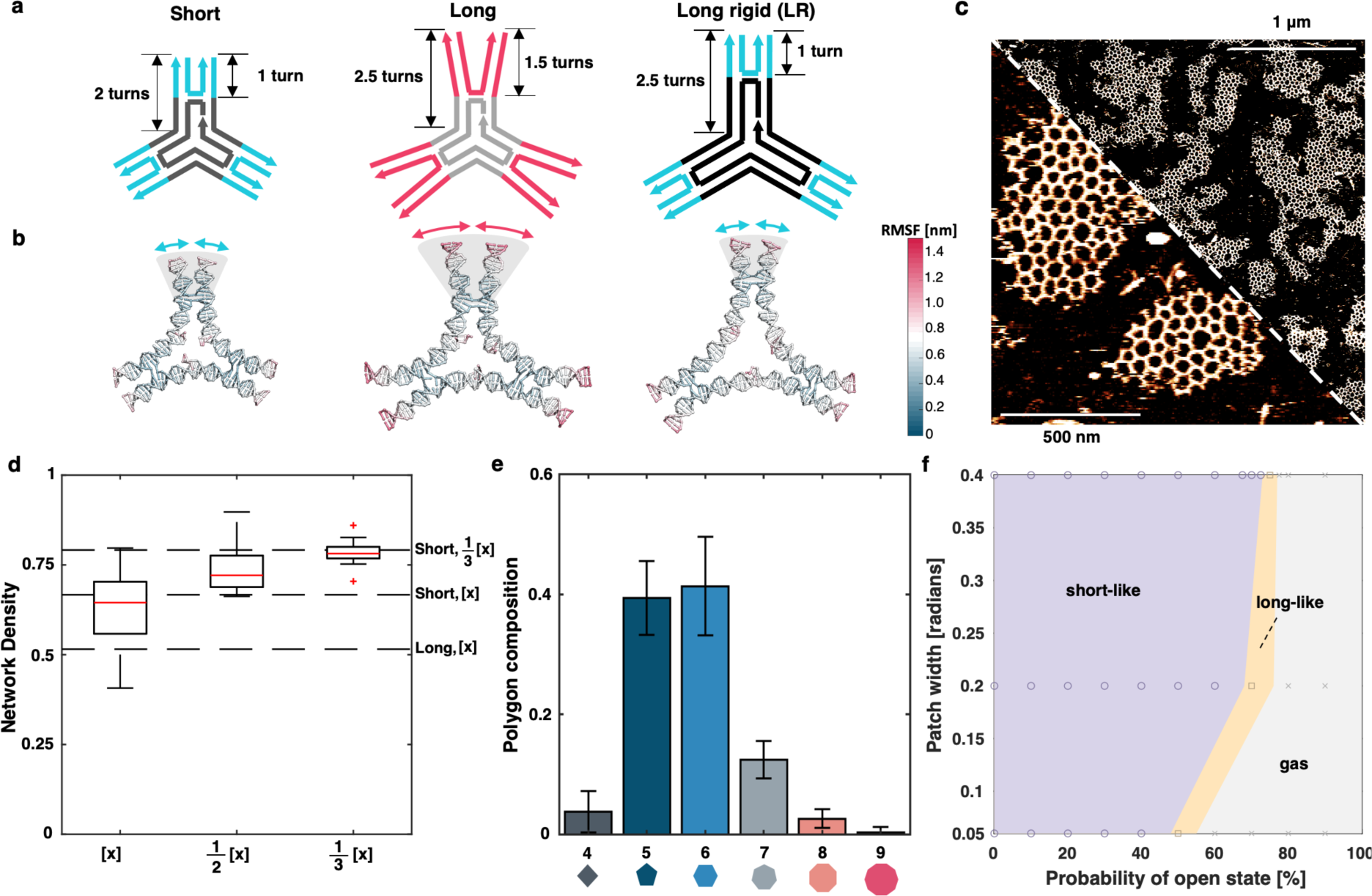
Restoration of peripheral rigidity. **a,** Design of “*long-rigid*” (LR) 3PS and comparison of local and global flexibility with the original *short* 3PS and *long* 3PS designs. **b,** representative frames from oxDNA simulations of the 3PS, coloured by the average RMSF. **c,** Steady-state AFM of LR 3PS, showing a restored radial large network assembly. **d,** ND of long rigid 3PS assemblies at decreasing concentrations ([x] = 3.8nM), dashed lines represent references for short rigid (SR) and long flexible (LF) monomers. The AFM images of 1/2x and 1/3x are shown in Supplementary Fig. 15. **e,** Polygon distributions for long-rigid 3PS assemblies, using at least 12 regions with an area of 750 x 750 nm. Error bars represent the standard deviation. **f,** Approximate phase diagram as a function of probability to be in the open state and the patch width for *σ=1*, *δ=0.038*. In purple *short*-like phase, in orange *long*-like phase, in grey gas phase. Each datapoint represents a set of conditions simulated.

We repeated all assembly experiments with the *long-rigid* 3PS motif. To our excitement, radially grown large networks were visible (Fig. 5c). Moreover, these were present in all dilutions with a network density score matching that of the *short* 3PS design (Fig. 5d). Interestingly, the polygonal composition seemed to contain more pentagonal defects as observed for the *short* 3PS monomers. Indeed, polygonal quantification confirmed the distribution to be more similar to that observed for the original *long* 3PS (Fig. 5e), however, with increased contributions of pentagons and higher-order polygons. Thus, while the peripheral rigidity restores the interface flexibility and mechanism of nucleation and growth, the crystallinity of these networks is still dominated by global flexibility.

Finally, we investigated whether the restoration of the radial growth could have resulted from a confounding factor such as the increased effective width of the interface resulting from the increased global flexibility. To do this, using our Patchy Particle model, we explored the parameter space described by the patch width, reflecting the global flexibility contribution to the interface, and the probability of being in an open state (Fig 5e). For the interaction strength defined by the regime of DNA blunt-end stacking, the dependency of the short-long phase boundary to the open/closed probability fell away with increasing patch width (value for 3PS particles is ∼0.3, based on the polygon fractions in Supplementary Fig. 14). This indicates that the assembly mechanism is indeed controlled by interface flexibility rather than global flexibility.

## CONCLUSION

In this study, we set out to explore the interplay between binding affinity and structural rigidity to control the growth and geometric order of supramolecular crystalline materials. To this end, we have exploited the programmable nature of the DNA macromolecule to engineer a series of monomers with closely similar shapes but nuanced differences in structural mechanics and chemical composition. Specifically, we used a tri-symmetric 3-point star (3PS), equipped with blunt-ends to promote assembly of polygonal networks on surfaces via directional *π*-*π* stacking. Initially, two designs were explored with *short* (2-turn) and *long* (2.5-turn) arms, respectively. For the *short* 3PS design, we observed the rapid nucleation and growth of predominantly hexagonal networks, while *long* 3PS failed to create structured networks. The minimal 5nt extensions in *long* 3PS introduced a global flexibility in the arms, as well as local flexibility at the 3PS periphery. This change in structural mechanics drastically altered the global self-assembly mechanism. HS-AFM allowed us to visualize and understand that the nucleation in the more flexible monomer was compromised, challenging stable growth of the supramolecular crystalline network.

The *π*-*π* stacking interaction has a high directionality, and thus a low spatial-tolerance. Consequently, small changes in orientation directly impact directional assembly. Nucleation seeds need to have a certain stability and lifetime to allow their transition into stable seeds and promote network growth, which is lost when the interface is too flexible. Using Monte-Carlo simulations, we demonstrated the critical effect of interface flexibility as the defining parameter of network formation. When the interface is too flexible, no stable nuclei can be formed and supramolecular-crystal formation is compromised. These observations are in line with earlier work on 3D DNA crystal formation, where gels constructed from too flexible monomers fail to crystalize.^52^ Only when interactions fall within the directional spatial tolerance of the non-covalent interaction engaged in the molecular design, the system can form stable nuclei and continue into a growth phase. When we restored the interface rigidity within the *long* 3PS monomers, the nucleation-growth mechanism was recovered. Spatial tolerance in directional multivalent interactions, classified as *interface flexibility*, comes forward as a critical parameter for the supramolecular self-assembly of macromonomers when long-range growth and order is desired.

Taken together, our study shows that a subtle change in local structural flexibility can have a dramatic effect on a systems’ global self-assembly, guided by interface flexibility. While demonstrated with DNA-based monomers in this study, these fundamental insights translate beyond the DNA helix, as any supramolecular network is assembled through non-covalent interfaces between macromonomers. Flexible-to-rigid transitions in biological macromonomers are surprisingly common, resulting from lateral interactions between monomers as seen in the example of the clathrin triskelion as well as via the addition of helper proteins in the case of the TRIM5a organization. With insights obtained in this study, we may start to look at biological transitions of flexibility and order with new appreciation for structural protein monomer design. Reversibly, the intentional introduction of interface flexibility can be used strategically when the formation of large supramolecular networks is not desired or should be disrupted (e.g. in the case of plaques, fibres, amyloids or even viral (dis)assembly). The presented study on the fundamental role of interface flexibility to control the mechanisms behind dynamic self-organization, enables the design of synthetic macromolecular building blocks showcasing predictable organization at the liquid-surface interface. Exploiting the global and local rigidity/binding affinity balance at biological interfaces presents new opportunities for the engineering of dynamic cellular nanotherapies and controlled growth -or disruption-of molecular networks.

## Methods

Full materials and methods and additional characterization can be found in the Supporting Information to this manuscript.

### Preparation of DNA 3PS motifs

For each 3PS, ssDNAs were mixed at the assigned molar ratios in the annealing buffer containing 5mM TRIS (Bio-Rad), 1mM EDTA (ITW Reagents) and 10mM MgAc_2_ (abcr GmbH) to give a final solution of 0.6 µM DNA motif in 50 µL. The DNA solutions were annealed at: 80 °C for 5 minutes, 60 **°**C for 10 minutes, cooled from 60 °C to 20 °C by 1 °C every 10 minutes, 20 °C for 10 minutes and stored at 4 °C.

### Static AFM imaging

40 µL of DNA tile at designated concentration was deposited on a freshly cleaved mica (grade V1, Ted Pella). Mica was placed in a petri dish containing 1mL water to create a humid environment and sealed with Parafilm. The sample was left overnight for self-organization on mica surface before imaging. All AFM images were acquired in tapping mode in liquid on a Cypher VRS (Asylum Research Inc.) using BioLever mini cantilever (BL-AC40TS-C2, Olympus).

### HS-AFM imaging

Images have been acquired on a home-built small cantilever AFM with photothermal excitation as described elsewhere.^35,36^ All images were acquired in photothermal off-resonance tapping mode at frequencies of 100kHz using AC10DS cantilevers (Olympus). 50µL of 10mM MgAc_2_ was injected close to the cantilever in the dedicated channel of the cantilever holder. The cantilever was approached to the surface and the surface was scanned to check for contaminations. The cantilever was retracted from the surface, the buffer extracted from the holder, and 50µL of the DNA 3-point-star (3PS) solution injected and imaged immediately. The sample was scanned at a line rate of 100 Hz (256 lines × 256 pixels) unless otherwise specified. The setpoint was kept at the lowest level required for proper tracking.

### Automated analyses of assembled networks

Extraction of information from AFM images has been carried through a custom MATLAB^30^ routine of (1) opening and standard AFM artefacts removal, (2) segmentation and skeletonization, (3) polygon and particle detection, and (4) collection of observables per connected component (island).

### OxDNA simulations

We prepared a vHelix^53^ in Maya^54^ model for all compounds with GC TA as terminal base couple, which was subsequently converted it in oxDNA^38,39^ compatible formats for coordinates and topology using TacoxDNA.^55^ Unless stated otherwise, simulations follow standard settings for the base dependent forcefield (oxDNA2) and a salt concentration of 0.5M. The pipeline involved a minimization of 1e6 steps using the steepest descent algorithm. 8 sets of molecular dynamics run in NVT ensemble where we progressively increased the temperature using the following ramp: 1, 1, 10, 10, 20, 20, 25, 25. Until this moment we kept the restraints between base-pairs using the standard settings generated with the related tool in OxView^56,57^ and used a timestep of 5e-4. We then set the timestep to 1e-3 and proceeded with a last simulation at 25 °C for 5e6 steps where we halved the strength associated to the restraints before starting our production run at the same temperature without base pair restraints. During the production runs, configurations were stored every 5e5 steps. Resulting files were converted to psf, pdb, and mdcrd files using an in-house python script. Analysis, snapshots and measures were then performed using VMD^58,59^ and MATLAB.

### Patchy-particle simulations

MonteCarlo simulations are run on a modified version of the engine developed by Hedges^60^, available online.^61^ A modified version of the Kern-Frenkel potential^45^ was implemented in order to take into account patch states and adhesion to mica. Detailed formulation of the potential and specific parameters used in each set of simulations are given in SI. The recorded trajectories were rendered as images using an in-house python script and VMD. The analysis of the frames was then performed in MATLAB following similar procedures as for the experimental AFM data (without step 1). Labelling of the states to construct the phase diagrams shown in Figure 4 and Figure 5 was done by evaluating absolute values and sudden drops in the MND values in function of the parameters of interest. Details about the potential used and the specifics of the simulations are reported in Supplementary Notes.

## Data availability

All data that support the conclusions in this manuscript are present in the main text or the supplementary materials. Source data are provided with this paper.

## Code availability

The simulation engine is available online at https://github.com/mosayebi/PatchyDisc. Analysis scripts for processing the trajectories are available upon reasonable request.

## Acknowledgments

M.M.C.B. and V.C. are grateful for the financial support by the Swiss National Science Foundation (SNSF), under the Eccellenza program (PCEGP2_181137). G.F. thanks the support from the H2020 - UE Framework Programme for Research & Innovation (2014-2020) and ERC-2017-CoG; InCell (773091), as well as the Swiss National Science Foundation through grant 200021_182562. M.M., T.B.L. and D.N.W. were supported by the UKRI-funded Synthetic Biology Research Centre, BrisSynBio (BB/L01386X/1). M.M.C.B., V.C and C.T. thank dr. Diana Morzy for critical comments and discussion in early states of the project.

## Notes

### Competing Interest Statement

The authors have declared no competing interest.

