## Supporting Information for "Structural flexibility dominates over binding strength for supramolecular crystallinity"

### Extended Methods

#### Oligonucleotides.

All DNA tiles designed in this study are based on the original 3-point-star (3PS) motif[1]. The modifications on the DNA sequences of the 3PS motif were made following the principles presented by Seeman[2]: All 4 nucleotide-long subsequences of individual DNA strands are (1) unique, (2) not self-complementary (e.g. TGCA) and (3) includes both purine and pyrimidine. The same rule applies for the 4 nt-long subsequences that are on a continuous DNA duplex but span a junction.

All oligonucleotides used in this study were either bought from Integrated DNA Technologies, Inc. or synthesized in-house on OligoMaker (TAG Copenhagen) following standard synthesis protocols. The sequences are listed in Table S1.

#### Purification of DNA tiles.

Preparation methods of individual DNA tiles can be found in the main text Materials and Methods. The annealed product was loaded on a 3% agarose (Sigma) gel and the gel was run at 60 V for 150 minutes in an ice-cooled water bath. The running buffer contained 0.5x TBE (Thermo Scientific, 44.5 mM Tris, 44.5 mM boric acid and 1mM EDTA; pH 8.0) and 10 mM $MgAc_2$. The band corresponding to the DNA motif was excised with a surgical blade and loaded in a Freeze 'N Squeeze gel extraction spin column (Bio-Rad). The column was centrifuged for 20 minutes at 3000 RCF and 4 $^{o}$C. To replace the buffer with the storing buffer (same with the annealing buffer), the flow through was collected and pipetted to a Vivaspin 500, MWCO 3000 (Sartorius). After an initial spin at 3000 RCF for 30 minutes, 300 $\mu$L of 1x storing buffer was added to the solution and centrifuged again at 3000 RCF for 90 minutes. This last step was repeated one more time to ensure that the buffer is replaced.

#### Native PAGE.

6% polyacrylamide gels were prepared following standard protocols. The gels were run at 120 V for 50 min in 0.5xTBE. The gels were stained with SYBR Gold (Sigma) and imaged with ChemiDoc MP (Bio-Rad).

**Further details on analyses of AFM images.**

After opening of an image through a third-party developed script[3] in MATLAB[4], the routine includes standard practices, such as: median line differences removal, adaptive thresholding to identify background, polynomial surface fitting of the background and median line removal using background as reference[5]. We iterated the process a second time after resizing the image to a standard pixel width of 0.34 nm. Finally, we applied a gaussian filter with standard deviation of 2 pixels and capped the final values between 2% and 98% of the values in the image.

For segmentation and skeletonization, a first enhancement of the contrast between foreground and background is obtained by applying a custom filter based on a kernel of radii equal to the size of the particle of interest and intensity that would reduce progressively from the center to the border, in a way that would take into account the different orientations that a particle can assume. Initial thresholding and average filtering of the background are then applied to further enhance the image. Afterwards, a preliminary skeletonization step based on the watershed transform is performed. Using the preliminary skeleton, we perform a second, more precise thresholding to obtain a mask. This mask is then further processed to enhance holes, remove full areas and fill scars. For the steady state images, when automatic filling of scars would fail due their excessive thickness, we would resort to manual correction of the original image and restart the process.

Finally, the mask is refined through morphological operators and a 4-connected skeleton is extracted through a standard skeletonization algorithm.

The polygon and particle detection part are based on the hypothesis that segments in an image connect the centers of two particles. Therefore, we proceed with the identification of segments in the skeletonized image considering that a segment is either: (1) shared by two polygons, (2) isolated or (3) forming an angle with another one. For each isolated connected component in the foreground, we try to divide it in segments based on principle 1 and then apply a Ramer-Douglas-Peucker algorithm (based on the native one implemented in MATLAB) in order to identify all the vertices in each segmented line. To facilitate the vertex detection algorithm, when possible, we define a list of initial candidate vertices, consisting of branched points and the point that is furthest away from the center of mass. Finally, each vertex is the center of a 3PS-particle and polygons can be identified by counting their vertices.

Finally, we check how many polygons and particles (and eventual properties like being in contact with the background) a connected component has and stores it in dedicated matrices.

**Detailed protocol for patchy-particle simulations.**

The current version of the engine used is made available online at https://github.com/mosayebi/PatchyDisc. Main modifications from the engine developed by Hedges[6] include: a dedicated routine to parse input conditions for each simulation from a json file, the definition of a new potential and variables, and implementations of moves to change the states of the patch.

The pairwise interaction between two particles is given by (see figure 4c)

$$U_{(r_{ij},\theta_i,\theta_j,p_i,p_j)} = \begin{cases} \infty, & r_{ij} < \sigma \\ -\varepsilon p_i p_j, & \sigma < r_{ij} < \sigma + \delta \wedge \theta_i < \theta_{pw} \wedge \theta_j < \theta_{pw} \\ 0, & otherwise \end{cases} \quad \text{Eq. S1}$$

where, $r_{ij}$ is the center-to-center distance between the two interacting particles, $i$ and $j$; $\sigma$ is the diameter of a particle; $\delta$ is the radial width for the interaction; the term $\varepsilon$ represent the patch-patch binding energy; $\theta_i$ is the smallest angle formed by the direction of any of the patches in particle $i$ and the line connecting the centers, similarly $\theta_j$ is the smallest angle formed by the direction of any of the patches in particle $j$ and line connecting the centers; $\theta_{pw}$ represents the maximum angle at which the interaction can happen; $p_i$ and $p_j$ are respectively the states of the aforementioned patches in $i$ and $j$ and they can either assume a value of 0 or a value of 1. This formulation corresponds to a standard Kern-Frenkel (KF) potential[7], with the extra condition that both patches only interact when they are in closed state.
In addition to the above modified KF potential, we consider a constant term ($W$) reducing the particle energy when at least one of its patches is involved in a patch-patch bond with another

particle. This term is motivated by our experimental observation that the interacting monomer engages with the mica surface better than an isolated monomer. As result, isolated monomers could not be resolved due to their high mobility, and yet dimers could be clearly observed in an AFM image.

$$H_{(r_{ij},\theta_i,\theta_j,p_i,p_j)} = U_{(r_{ij},\theta_i,\theta_j,p_i,p_j)} + W \qquad \text{Eq. S2}$$

The moves in our Monte Carlo (MC) simulation are either translation/rotation of a particle or changing the state of a single patch (which is attempted 3 times more often than the translation/rotation move). To ensure detailed balance, we accept moves according to the Metropolis rule, where the energy change between the original configuration "$a$" and the resulting configuration "$b$" is calculated as

$$\Delta G_{a\to b} = \Delta H_{a\to b} + \Delta P_{a\to b}, \qquad \text{Eq. S3}$$

where $\Delta H_{a\to b}$ represents the energy change between the two configurations based on the potential defined in Eq.S1, $\Delta P_{a\to b}$ is the free energy change resulting from switching the patch state while moving from $a$ to $b$, based on the probability of the patch being in "open" state, $P_o$, when not interacting with others (in our case, for a particle $i$ in a simulations of $N$ partilces, $\sum_{0,j\neq i}^{N} U_{ij} = 0$). Since we have only 2 possible states the specific formulation of the second term is

$$\Delta P_{a\to b} = \begin{cases} 0, & p_a = p_b \\ \log\left(\frac{P_o}{1-P_o}\right), & p_a = 0 \wedge p_b = 1 \\ \log\left(\frac{1-P_o}{P_o}\right), & p_a = 1 \wedge p_b = 0 \end{cases} \qquad \text{Eq. S4}$$

Where $p_a$ represent the state of the patch in state $a$ and $p_b$ represent the state of the patch in state $b$.

The units of length in the simulations are normalized using as reference length the diameter of a particle (16 nm in our case), the energy is normalized by $k\,T$, where $k$ is Boltzmann's constant, $T$ is the thermodynamic temperature and angles are measured in radians.

Simulations are performed in 2D placing 1500 particles over an area of 150x150 normalized units (~1.6x1.6 $\mu m^2$), matching densities measured for our low-density images. The total number of steps is 2.4e7 for each simulation. The interaction width is set at 0.038, implying that a π-π stacking would have a width of ~0.6 nm[8]. Respecting the conditions of the one bond per patch regime[8], $\theta_{pw}$ has been tested between 0.06 and 0.40 radians (used for Fig. 5). $\varepsilon$ has been tested between 2 to 10 (used for Fig 4). In Fig 5, $\varepsilon$ was kept at 6. $W$, when a particle is interacting with another particle, has a value of 6, otherwise 0. Particle positions and orientations are initialized randomly, while each patch state is initialized to 0.

**Specifications for simulations of isolated monomers in solutions.**

The protocol followed the one reported in Materials and Methods. Production runs consisted of 1.85e8 steps. 9 replicas were performed for each monomer, respectively: short, long and long rigid. During analysis we discard the first 1e7 steps and considered frames collected every 5e5 steps. In Figure S10 are reported for each compound characteristic configurations and the RMSF computed over all the trajectories.

**Specifications for simulations of isolated monomers on surface.**

In a similar fashion to what has been done in Materials and Methods, we prepared flat *de novo* designs of our monomers. However, these models have been used to select the starting configurations from the trajectories simulated in the previous section as follows: for each monomer, we selected as starting configuration the one that would have the lower RMSD compared to the manually constructed flat models. Afterwards, we manually aligned the starting structures to a plane perpendicular to the y axis using OxView[9,10], and then applied the rest of the protocol described in Materials and Methods. In this context, we run simulations while implementing two repulsion planes, confining the compound in a layer of around 2 nm, starting with a force constant of 0.3 for the minimization. Additionally, we increased the steps in the minimization process to 7.5e6. Throughout the equilibration NVT simulations we slowly increased the force of the repulsion plane, by setting the associated parameter to: 0.3, 0.6, 0.9, 1.2, 1.5, 1.8 and 2.1. The last was then kept throughout the last equilibration run and production.

Production runs consisted in 1.35e8 steps. 12 replicas were performed for each monomer. During analysis we would discard the first 1e7 steps and considered frames collected every 5e5 steps.
In Fig. S11 are reported for each compound characteristic configurations and the RMSF computed over all the trajectories; end to end distance are plotted in Fig. S13.

### Supplementary Figures and Tables

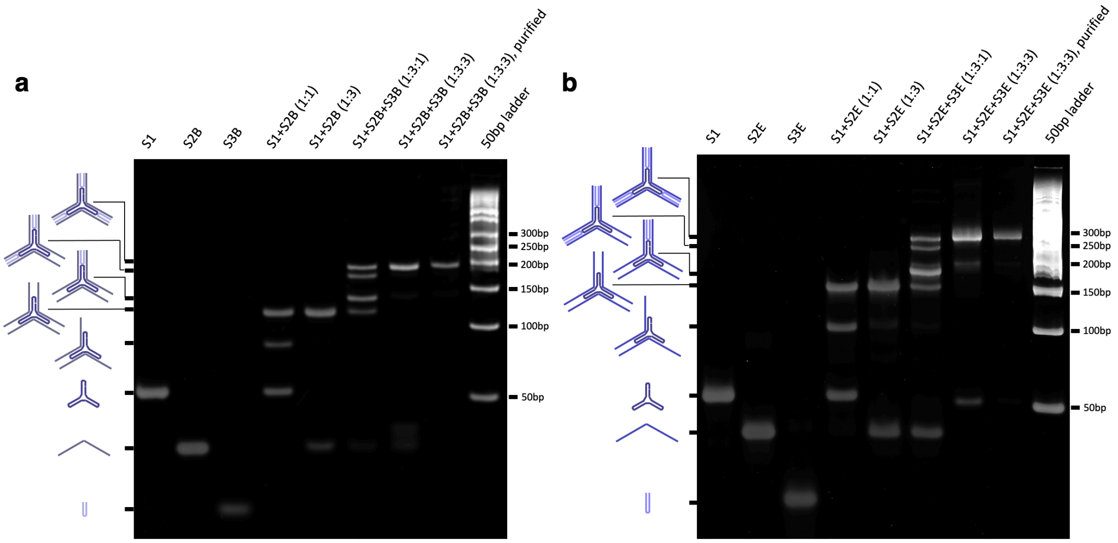


**Fig. S1. Native PAGE (6%) analysis of the formation of (a) short GCTA and (b) long GCTA DNA 3PS.**

The compositions of the samples and the structures corresponding to each band are shown above and on left of the gels respectively.

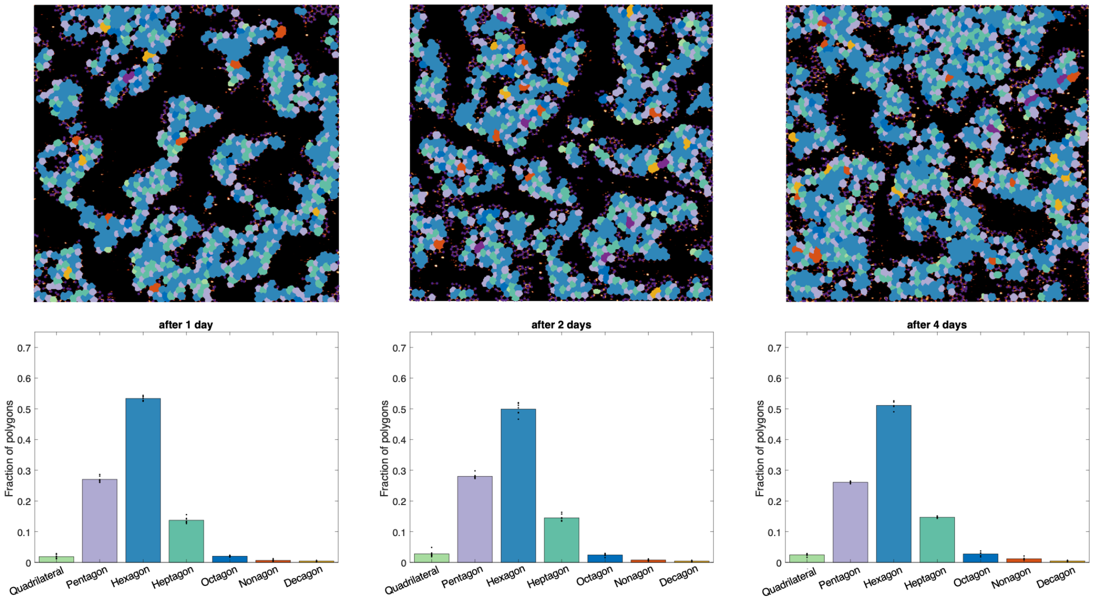


**Fig. S2. The evolution of the assembly after 1-4 days.**

Bars present a weighted mean based on the number of monomers observed in an image. Each black dot represents the fraction of a polygon in a unique 750x750 μm$^2$ area. After a day of incubation, the distribution of polygons remains constant, prompting us to opt for a one-day incubation period on mica. While the surface coverage continues to increase gradually due to the slow evaporation of the sample, the assembly properties remain unaltered.

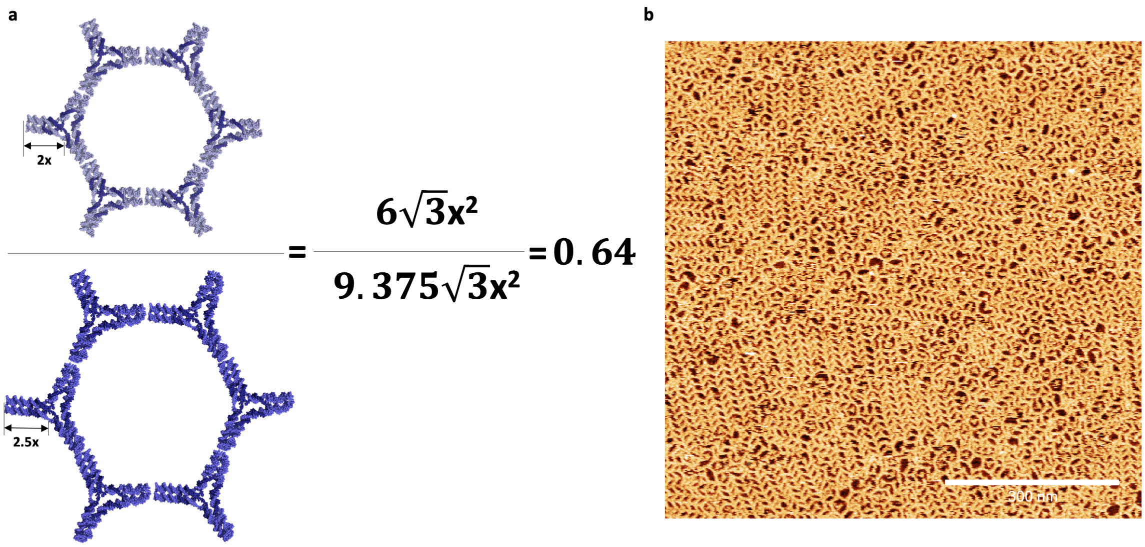


**Fig. S3. DNA concentration for AFM experiments.**

**a,** Long 3PS concentration. **x** indicates a turn of dsDNA. To cover the surface as much as short 3PS, 0.64 x 6nM = 3.84nM of long 3PS was used in the experiments. **b,** Overcrowding. In the cases of excessively high concentration, some blunt-end interactions are disrupted and the 3PS pack in a denser final form that minimizes the void between them.

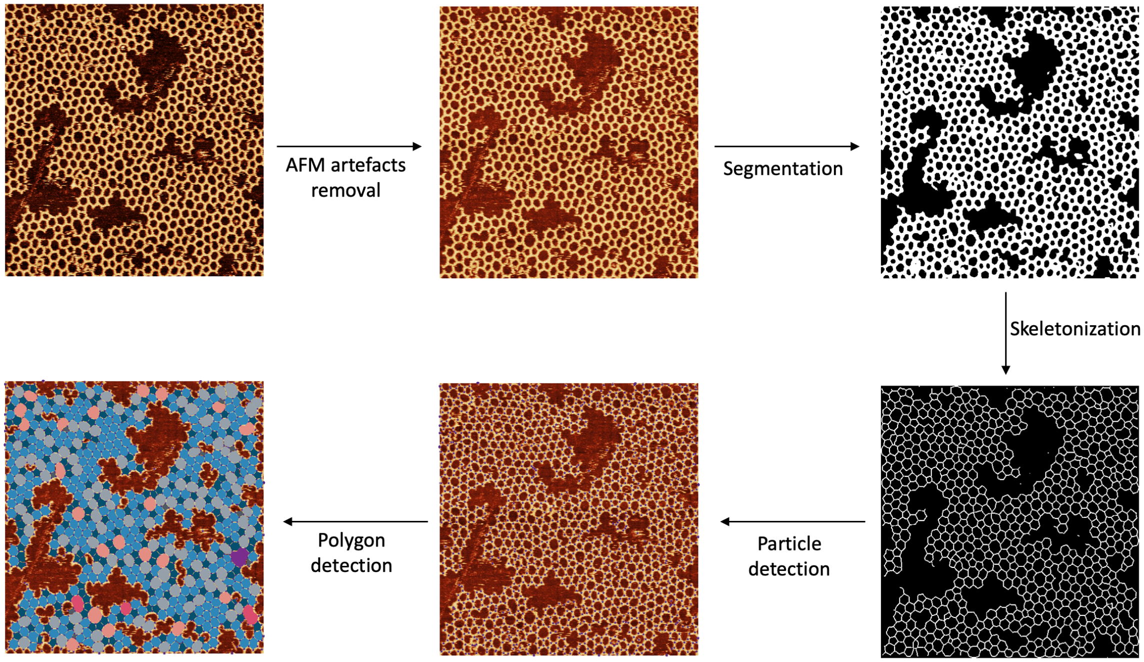


**Fig. S4. Image processing and analysis flowchart**

The typical image processing and analysis workflow includes (1) AFM artefact removal, (2) segmentation, (3) skeletonization, (4) particle detection, and (5) polygon detection.

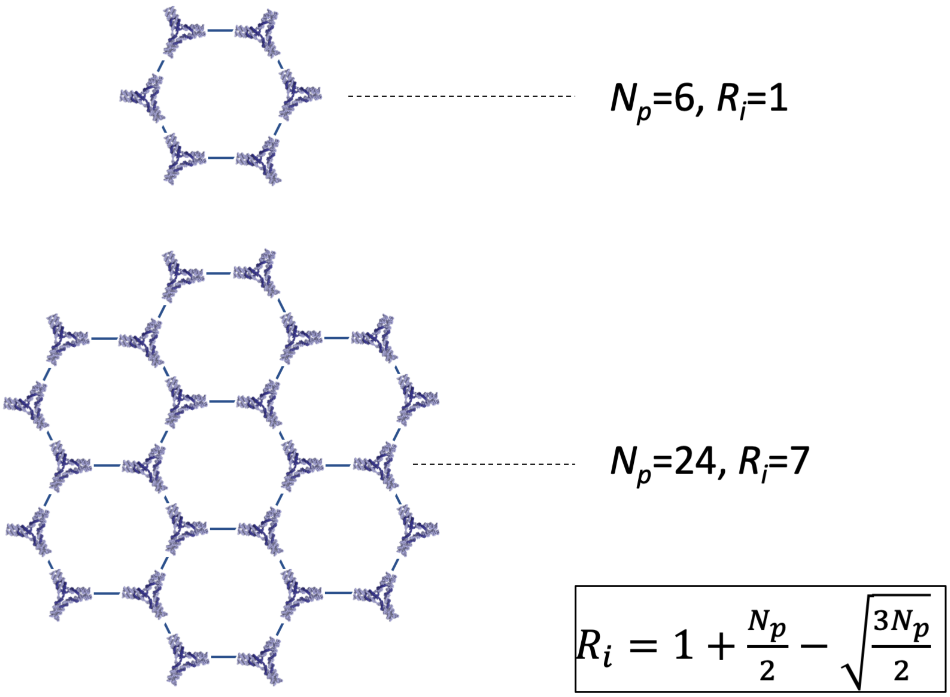


**Fig. S5. Ideal island for the calculation of Network Density (ND).**

ND is the ratio between the number of rings present in an island and the number of rings in an ideal island ($R_i$) formed by an equivalent number of particles ($N_p$). The term "ideal island" refers to a flawlessly radial structure devoid of any defects.

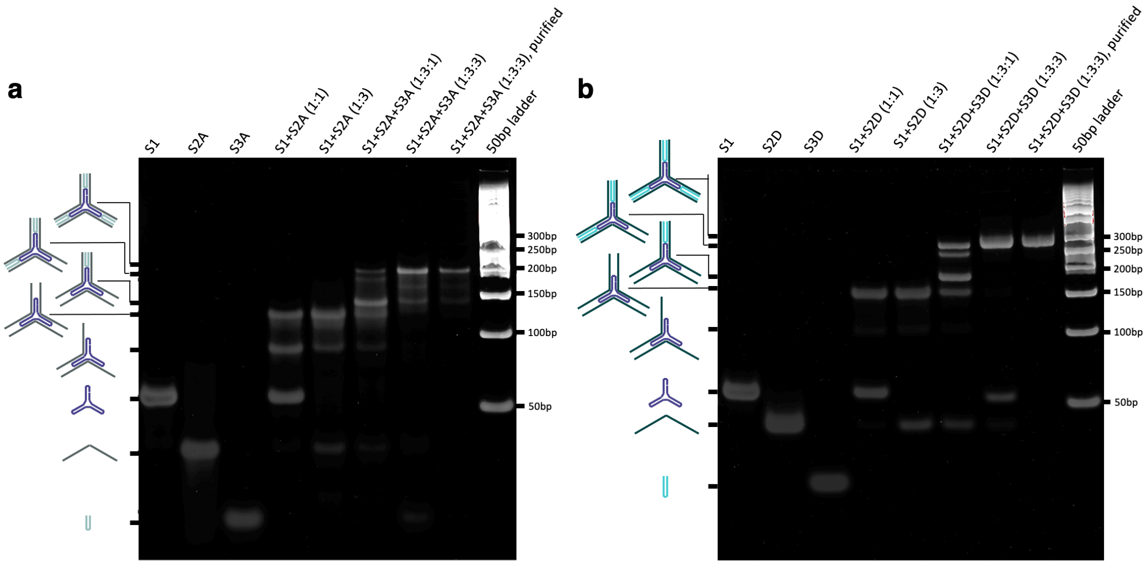


**Fig. S6. Native PAGE (6%) analysis of the formation of (a) short TATA and (b) long TATA 3PS.**

The compositions of the samples and the structures corresponding to each band are shown above and on left of the gels respectively.

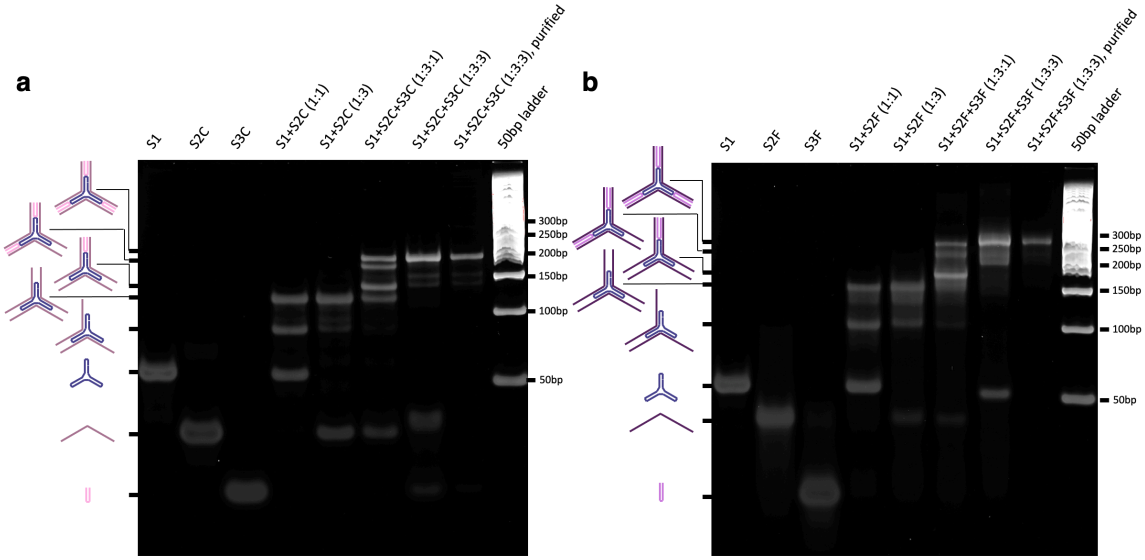


**Fig. S7. Native PAGE (6%) analysis of the formation of (a) short GCGC and (b) long GCGC 3PS.**

The compositions of the samples and the structures corresponding to each band are shown above and on left of the gels respectively.

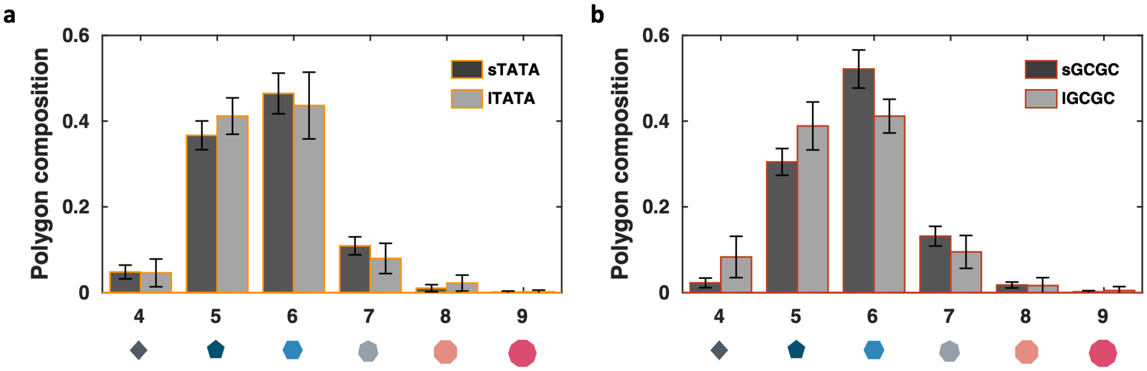


**Fig. S8. Polygon compositions at the steady state for (a) TATA and (b) GCGC.**

The polygon composition of TATA and GCGC 3PS after a day of incubation on mica. The bars include the data of at least 12 images (750x750 μm$^2$) per condition and they present a weighted mean based on the monomers observed in an image. Error bars indicates the standard deviation.

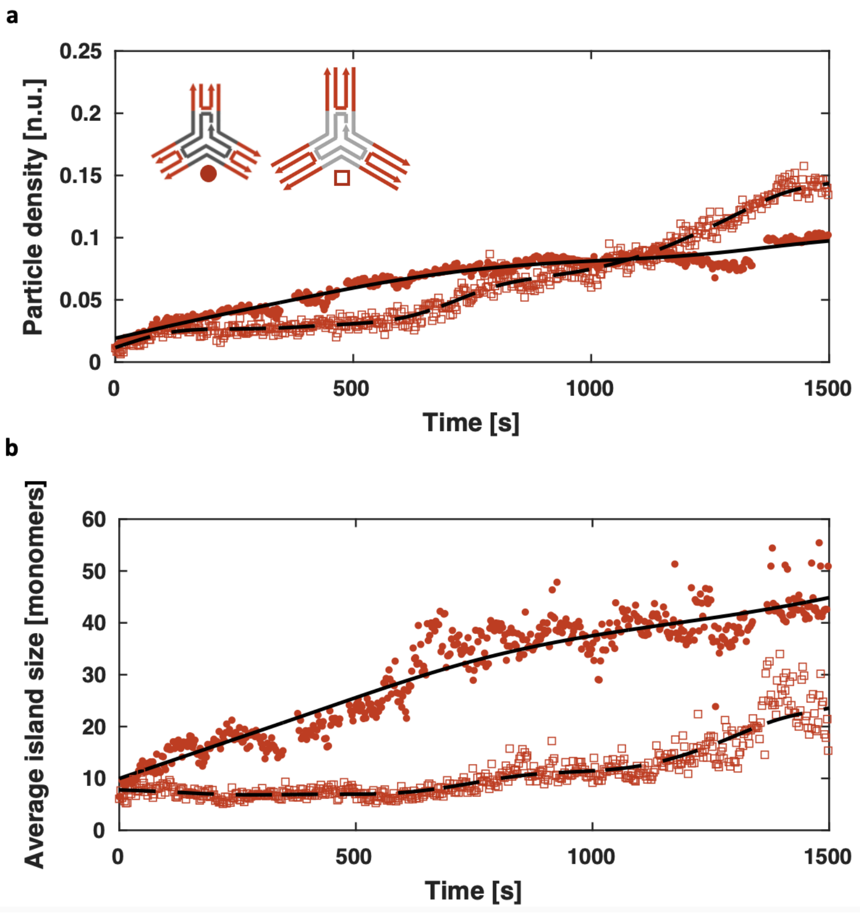


**Fig. S9. Dynamic self-assembly of short and long GCGC.**

**a**, Particle density on mica as a function of time. The number of particles on surface monotonically increases for both long and short 3PS. **b**, Average island size as a function of time. The islands formed by the short 3PS continuously increases with the arrival of more monomers on the surface. This strongly supports the growth phase of the assembly mechanisms, with every new monomer attaching to an existing island. Contrary, the islands formed by the long 3PS lack any kind of growth, meaning all incoming monomers try to make new nuclei. This changes around 1000s, where the surface density is so high, the small islands start to coalesce.

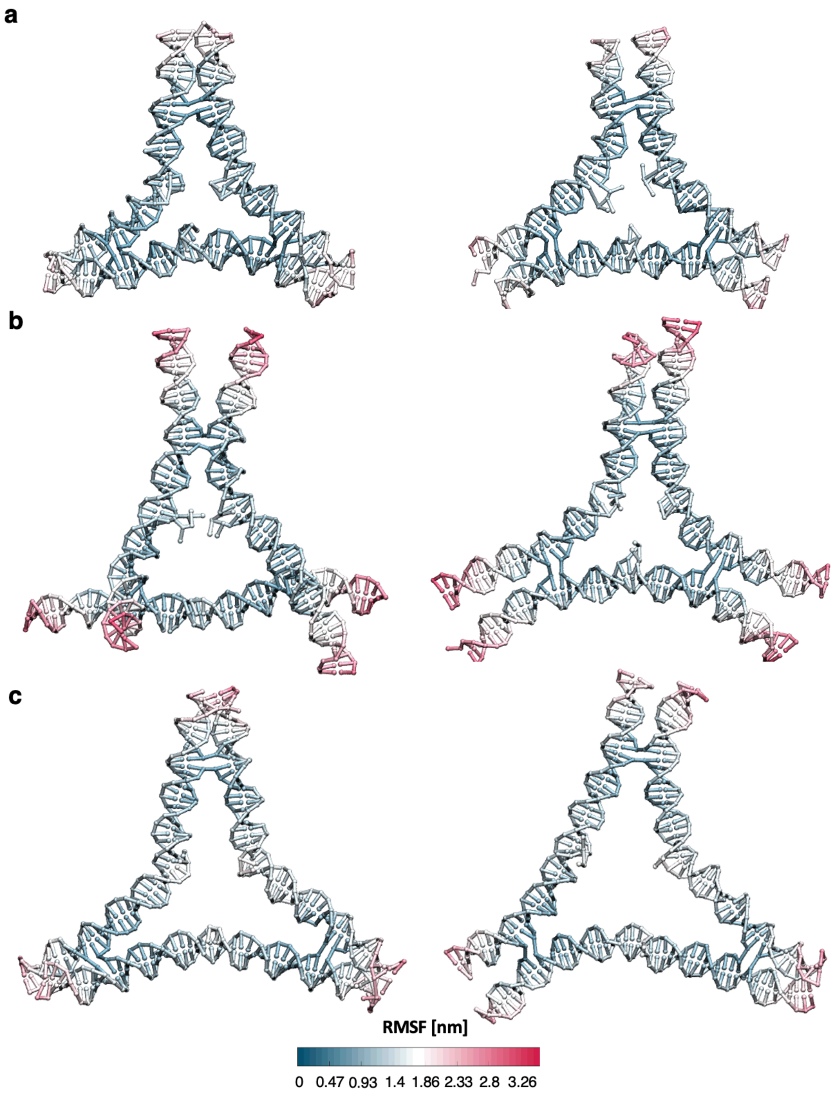


**Fig. S10. OxDNA simulation frames of (a) short, (b) long and (c) long rigid DNA 3PS in solution.**

The left frames display the conformations with the lowest RMSD when compared to the average conformation. The right frames depict the conformations closest to the flat ones.

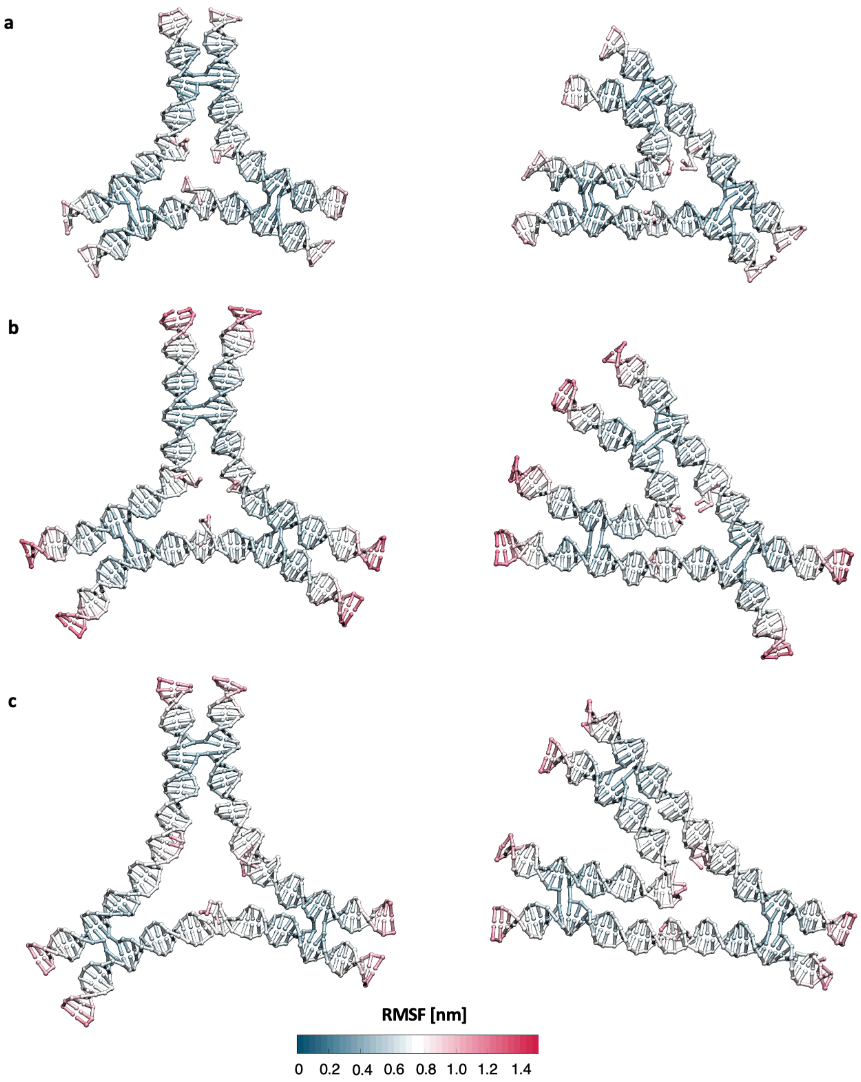


**Fig. S11. OxDNA simulation frames of (a) short, (b) long and (c) long rigid DNA 3PS confined on the surface.**

The left frames display the conformations with the lowest Root Mean Square Deviation (RMSD) when compared to the average conformation. The right frames depict the conformations with the highest RMSD which are less likely to be observed.

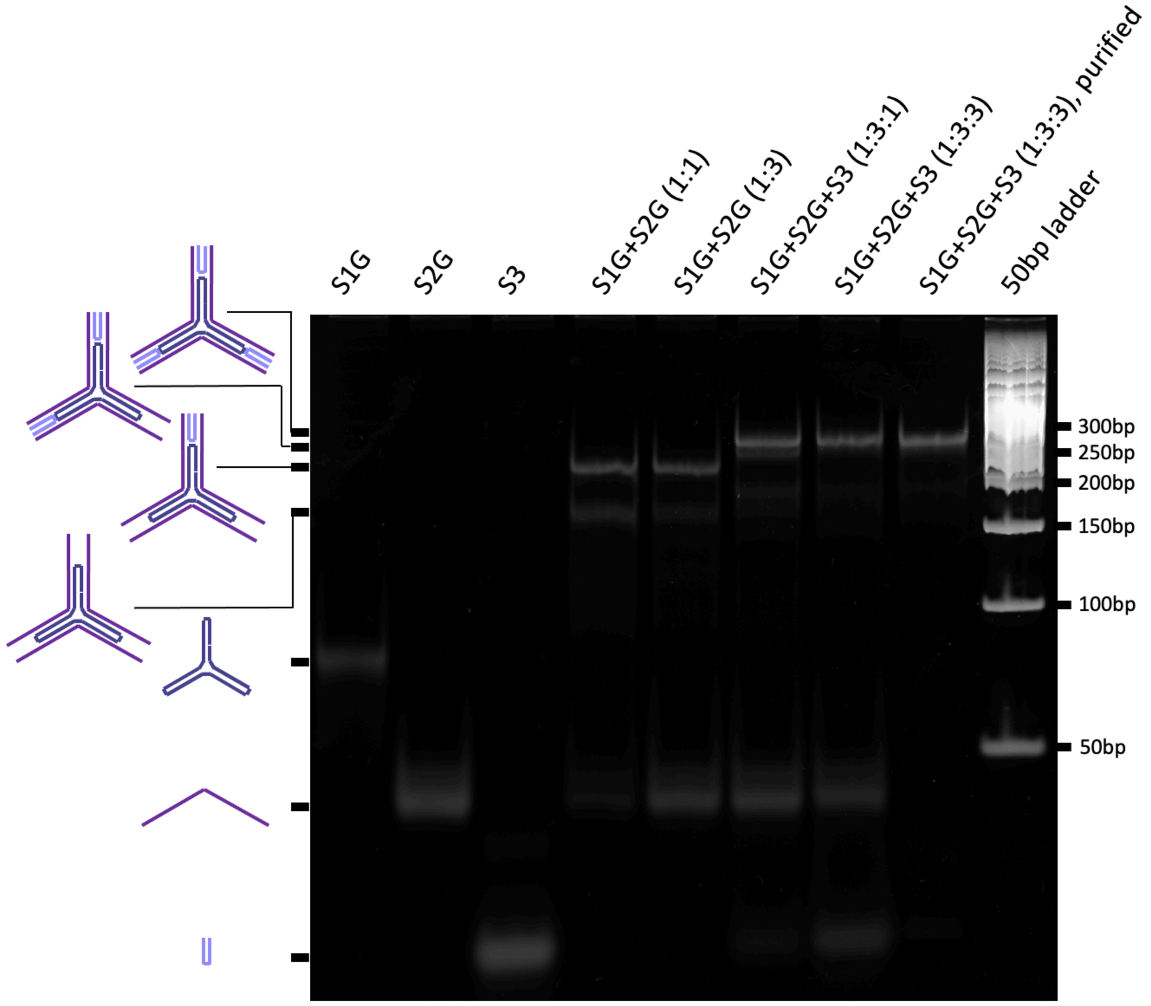


**Fig. S12. Native PAGE (6%) analysis of the formation of long rigid (LR) 3PS.**
The compositions of the samples and the structures corresponding to each band are shown above and on left of the gel respectively.

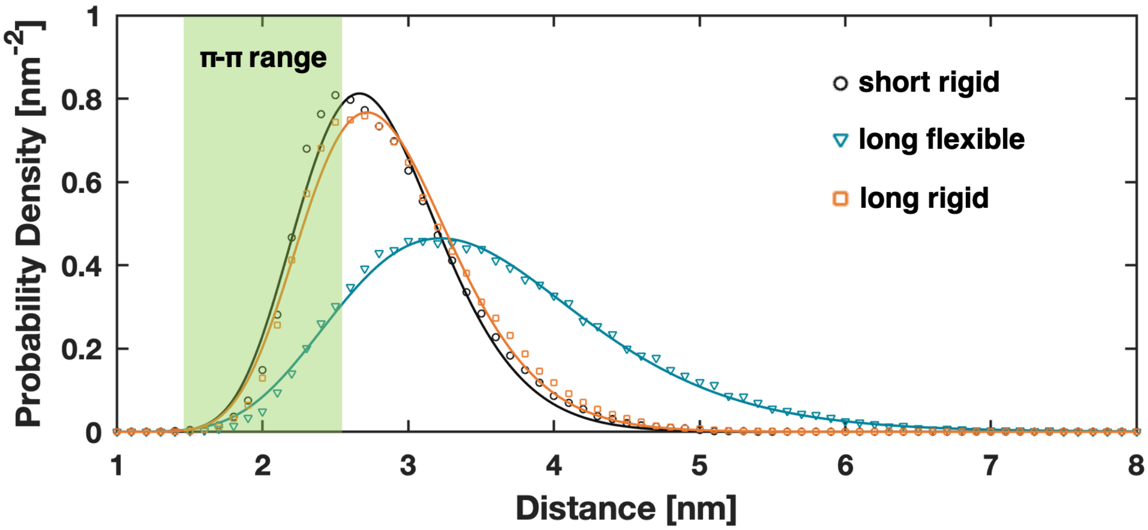


**Fig. S13. Probability distribution function of 3PS end-to-end distances based on OxDNA simulations.**

Each 3PS was forced to be in a plane as explained in the Extended Methods. The data for the short and long are already reported in Fig. 4b. Long rigid 3PS having a similar end-to-end distance distribution to short 3PS is a strong indication that their interface flexibility is similar.

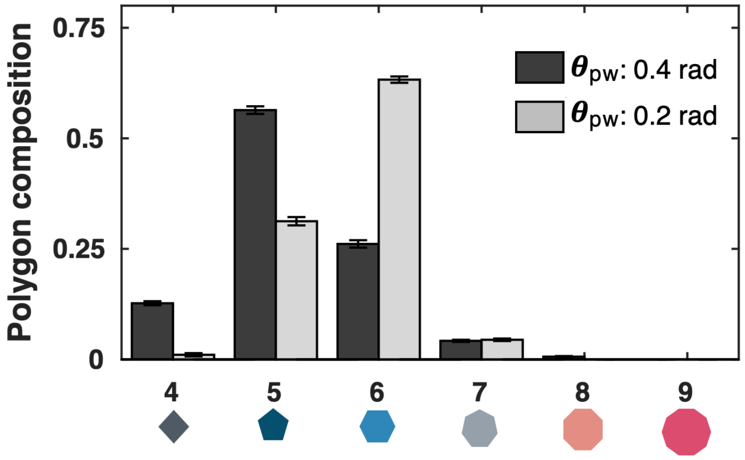


**Fig. S14. Polygon composition in the patchy particle simulations.**

In this set of simulations $\varepsilon$ , $P_o$ were 6 and 0.3 respectively. Since the fraction of hexagons we measure *in vitro* is ~0.5 which is included in the interval observed *in silico* (0.24<x<0.67), we expect the patch widths that correctly represent our constructs to be bound between 0.2 and 0.4 radians.

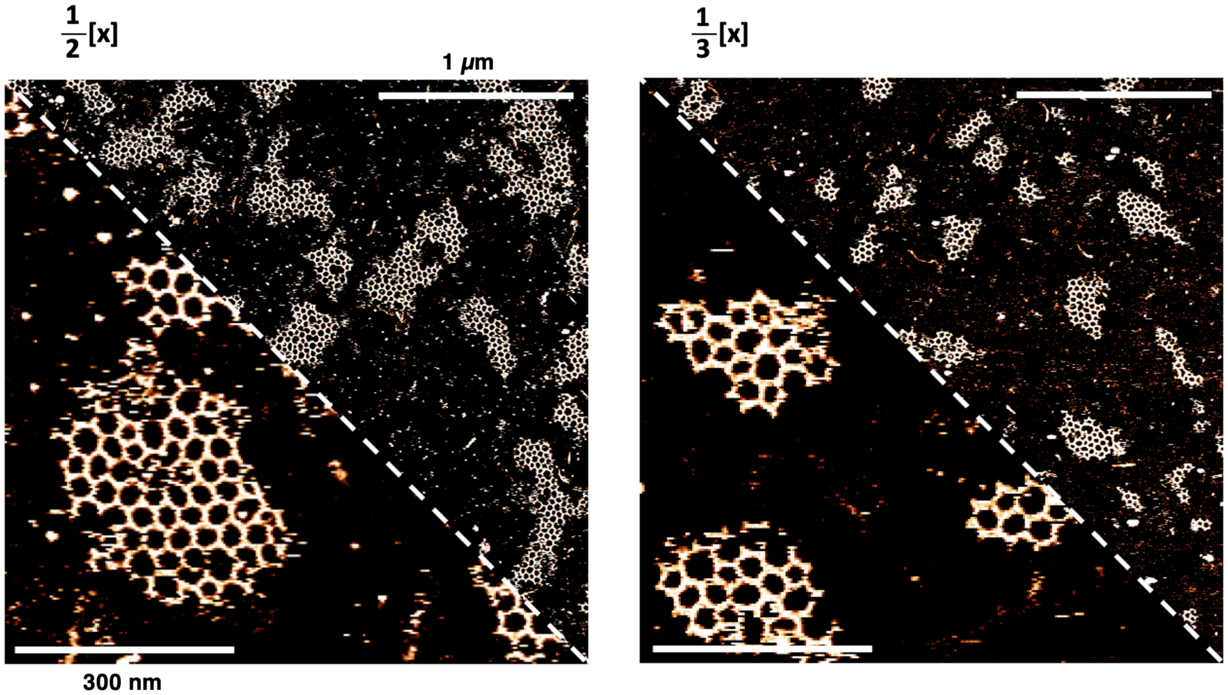


**Fig. S15. AFM images of the long rigid 3PS self-assembly at lower concentrations**
Long rigid 3PS assembles into radial islands at lower concentrations demonstrating its similar assembly mechanisms as short 3PS.

**Table S1. DNA motifs and the base sequences of the strands.**

<table>
<tr><th>Motif</th><th>Strand</th><th>Sequence</th></tr>
<tr><td>Present in all motifs except long rigid</td><td>1</td><td>5’-AGGCACCATCGTAGGTTTCTTGC<br>CAGGCACCATCGTAGGTTTCTTGCC<br>AGGCACCATCGTAGGTTTCTTGCC-3’</td></tr>
<tr><td rowspan="2">sTATA<br>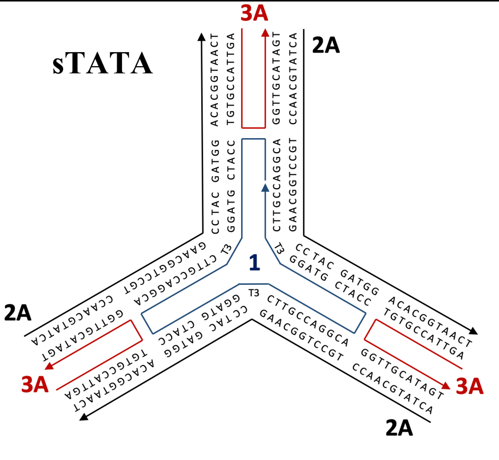</td><td>2A</td><td>5’-ACTATGCAACCTGCCTGGCAAGC<br>CTACGATGGACACGGTAACT-3’</td></tr>
<tr><td>3A</td><td>5’-AGTTACCGTGTGGTTGCATAGT-3’</td></tr>
<tr><td rowspan="2">sGCTA<br>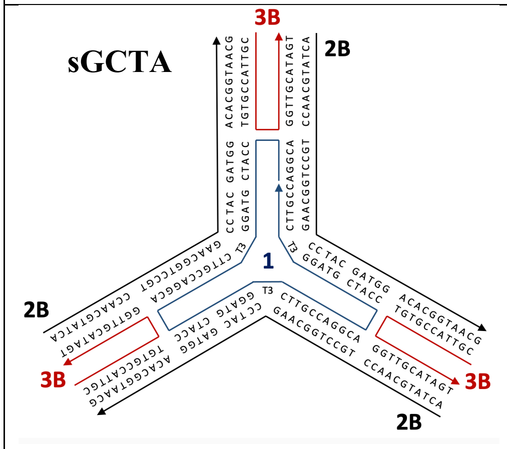</td><td>2B</td><td>5’-ACTATGCAACCTGCCTGGCAAGC<br>CTACGATGGACACGGTAACG-3’</td></tr>
<tr><td>3B</td><td>5’-CGTTACCGTGTGGTTGCATAGT-3’</td></tr>
<tr><td rowspan="2">sGCGC<br>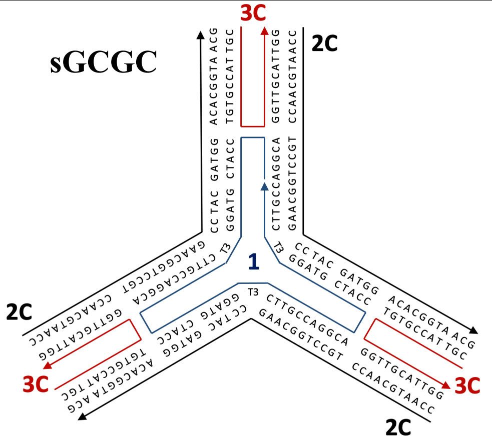</td><td>2C</td><td>5’-CCAATGCAACCTGCCTGGCAAGC<br>CTACGATGGACACGGTAACG-3’</td></tr>
<tr><td>3C</td><td>5’-CGTTACCGTGTGGTTGCATTGG-3’</td></tr>
</table>

| | | |
|---|---|---|
| ITATA 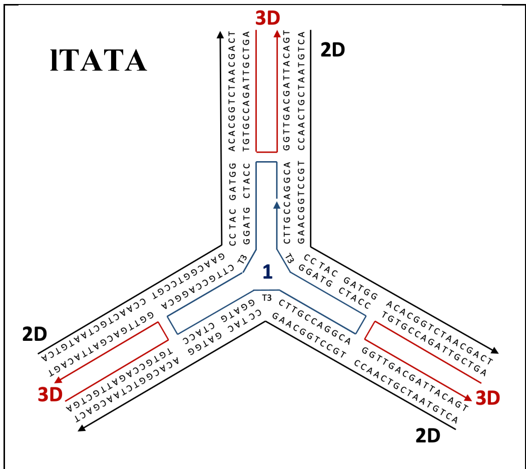  | 2D | 5'-ACTGTAATCGTCAACCTGCCTGGC AAGCCTACGATGGACACGGTCTAAC GACT-3' |
| | 3D | 5'-AGTCGTTAGACCGTGTGGTTGAC GATTACAGT-3' |
| IGCTA 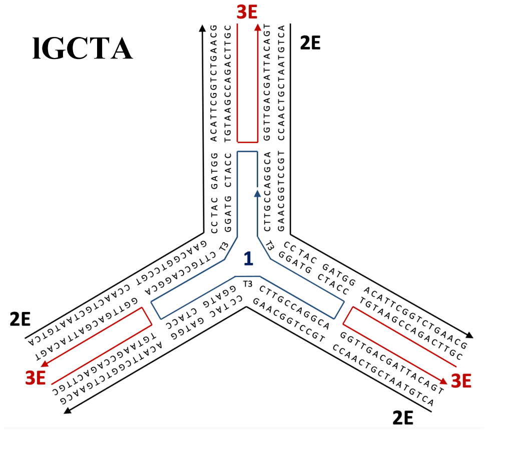  | 2E | 5'-ACTGTAATCGTCAACCTGCCTGGC AAGCCTACGATGGACATTCGGTCTG AACG-3' |
| | 3E | 5'-CGTTCAGACCGAATGTGGTTGA CGATTACAGT-3' |
| IGCGC 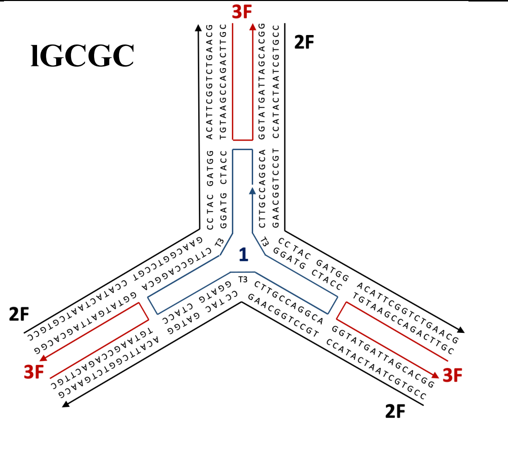  | 2F | 5'-CCGTGCTAATCATACCTGCCTGGC AAGCCTACGATGGACATTCGGTCTG AACG-3' |
| | 3F | 5'-CGTTCAGACCGAATGTGGTATGATT AGCACGG-3' |

| | | |
|---|---|---|
| Long rigid (LR) 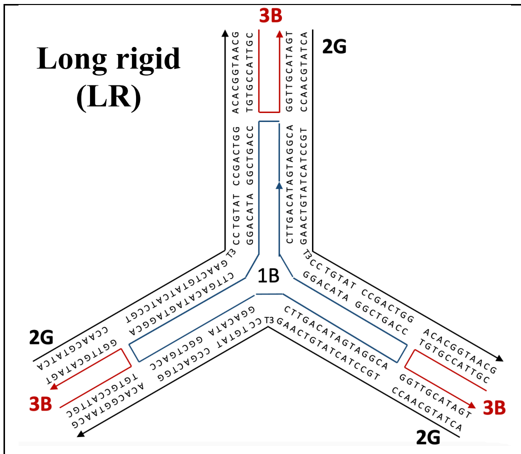  | 1B | 5'-AGTAGGCACCAGTCGGATACAGG CTTGACATAGTAGGCACCAGTCGGA TACAGGCTTGACATAGTAGGCACCA GTCGGATACAGGCTTGACAT-3' |
| | 2G | 5'-ACTATGCAACCTGCCTACTATGTC AAGTTTCCTGTATCCGACTGGACACG GTAACG-3' |

### Supplementary Movies

**Movie S1. Dynamic self-assembly of short GCGC**

**Movie S2. Dynamic self-assembly of long GCGC**

**Movie S3. Dynamic self-assembly of short GCTA**

**Movie S4. Dynamic self-assembly of long GCTA**

**Movie S5. Dynamic self-assembly of short TATA**

**Movie S6. Dynamic self-assembly of long TATA**

**Movie S7. Patchy-particle simulation of short**

**Movie S8. Patchy-particle simulation of long**
